# Multiscale chromatin modeling of chromosome X structural changes upon inactivation highlights the differential regulatory mechanism of *Xist*

**DOI:** 10.64898/2026.01.20.700676

**Authors:** Sangram Kadam, Tamar Schlick

## Abstract

The fundamental process of X-chromosome inactivation (XCI) involves silencing one X chromosome in female mammals by the *Xist* gene within the X-inactivation center (*Xic*). While experiments have identified key regulatory elements controlling *Xist* expression, mechanistic details are unknown. By combining nucleosome-resolution and coarse-grained polymer modeling, we reveal multiscale *Xic* reorganization during XCI driven by loop extrusion and epigenetic modifications, including methylation. At the nucleosome level, inactive X shows differential gene organization, where *Xite* is buried and compacted but *Xist* folds into a fragmented open structure; clutch patterns also change upon inactivation, and changes in methylation (in *Xite*) and NFRs (in *Xist*) explain the reorganization. At the megabase scale, our simulations reveal spatial rewiring: *Linx* -*Tsix* contacts are disrupted, isolating *Tsix* from its activator, while *Xist*, *Jpx*, and *Ftx* coalesce into an active compartment. This hierarchical and differential reorganization creates a chromatin architecture with an active domain for *Xist* favoring its expression, while a repressed *Xite* prevents *Tsix* reactivation. These general principles of how 3D genome organization directs development have implications for diseases related to XCI and extend beyond XCI to gene regulation broadly.

## INTRODUCTION

X-chromosome inactivation (XCI) is a fundamental epigenetic process by which one of the two X chromosomes in female mammals is transcriptionally silenced. This important process ensures proper dosage of X-linked genes between female XX chromosomes and male XY chromosomes [1–3]. The XCI begins when the *Xist* gene in the X-inactivation center (*Xic*) is expressed on only one of the two X chromosomes (Fig. 1). The long non-coding RNA transcript of *Xist* then spreads across the entire X chromosome and recruits chromatin-modifying complexes that condense the chromosome into an inactive structure [4–7]. How this mechanism of asymmetric expression of *Xist* on only one chromosome leads to chromosome-wide epigenetic repression is an important challenge in developmental biology [8–11]. It also has broad implications for understanding XCI-related diseases such as Rett, fragile X, Fabry disease, systemic lupus erythematosus, and the development of therapies for them [12–16].

**FIG. 1.**
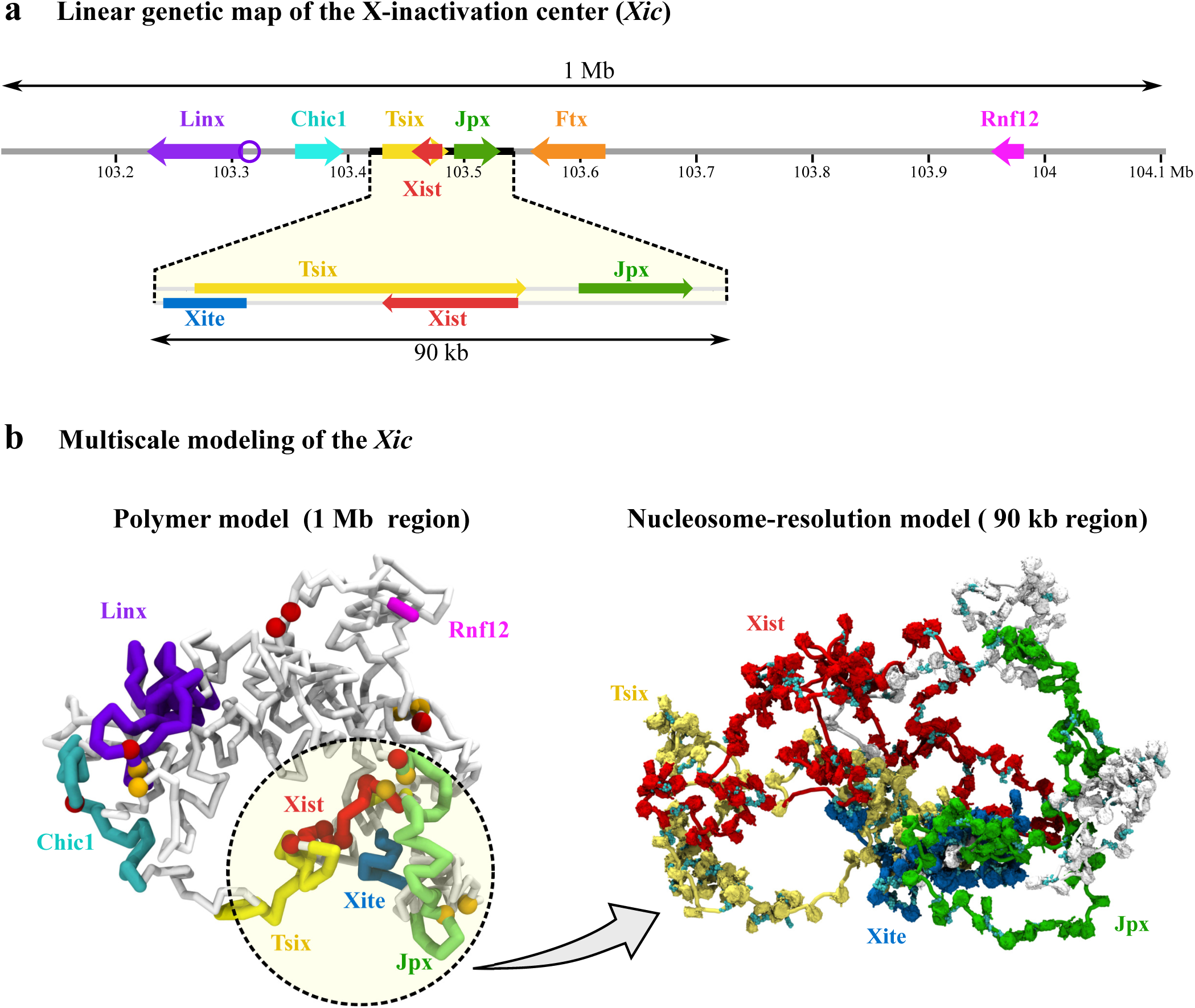
Multiscale modeling strategy for the X inactivation center, which combines a nucleosome-resolution mesoscale model (90 kb region) with a polymer model of the extended 1Mb *Xic* region. **a** Genes and regulatory elements across the 1 Mb region are annotated above, with a zoomed-in view of the 90 kb segment—encompassing *Xite*, *Tsix*, *Xist*, and *Jpx* —shown below. **b** We use a multiscale modeling approach to model *Xic*. Chromatin reorganization of the *Xic* region during XCI is modeled by a coarse-grained polymer model with loop extrusion (2kb resolution) to investigate interactions between genes that are hundreds of kilobases apart; this is coupled to a nucleosome-resolution modeling (9bp resolution) to examine gene organization at the scale of 90kb. The red and orange beads in the polymer model denote CTCF sites and the anchor sites of bound cohesins.

Key in initiating, establishing, and maintaining the silenced state on the X chromosome is the Xic locus [17, 18]. Besides the *Xist* gene, other Xic genes and regulatory sequences play an important role during XCI [5, 19]. Together, these genes form a multiscale regulatory network controlling the expression of *Xist* through intricate three-dimensional (3D) interactions (Fig. 1). We know that the precise coordination among these sequences ensures that *Xist* is expressed on one X chromosome, which becomes inactive (Xi), while the other remains active (Xa). However, the spatial organization of these regulatory elements and how their 3D organization influences gene expression and RNA spreading remains poorly understood. Understanding the complex interplay between these regulatory sequences and the accompanying dynamic changes in chromatin organization within the *Xic* region is crucial for deciphering the molecular basis of XCI and addressing related human diseases.

Many experimental studies have explored the *Xic*’s genome organization and how its regulatory sequences interact with each other to control the expression of *Xist*. In mouse embryonic stem cells, *Xic* is organized into two TADs, with *Tsix*, a negative regulator of *Xist*, present in the opposite TAD from that of *Xist* [18, 20]. The complex network of positive and negative regulators in the *Xic* region includes *Tsix*, *Jpx*, *Ftx*, *Rnf12*, *Xite*, and *Linx* (Fig. 1). All compete to control the expression of *Xist* to determine the transformation into Xa and Xi [21–30]. *Tsix* and its positive regulator *Xite* prevent *Xist* accumulation on Xa and are crucial for X-chromosome selection [18, 27, 28, 31, 32]. In addition to these genes, *Jpx*, *Ftx*, and *Rnf12* are thought to upregulate *Xist* (i.e., increase its expression) [21–26, 33], while the *Linx* promoter is known to downregulate *Xist* through direct contact *in cis* [29, 30]. Although we know that these regulatory sequences are crucial for controlling the expression of *Xist*, a key question remains: how do the physical interactions among these regulatory sequences influence the gene expression of *Xist*Evidence is accumulating from chromatin conformation capture experiments that such genome organization is important to XCI on several spatial levels [18, 34].

Indeed, the formation and maintenance of TADs themselves present another layer of complexity. Experimental evidence shows that, while cohesin-mediated loop extrusion is crucial for TAD formation on active X, inactive X shows two megadomains and weakened TAD structures [34–39]. Moreover, experiments reveal that *Xist* RNA evicts cohesin from Xi, while *Jpx* RNA regulates the selection of CTCF binding sites through competitive inhibition [6, 40]. This suggests that modulation of CTCF-cohesin loop extrusion through these different RNAs provides an additional pathway to reorganize chromatin structure during X-inactivation. The molecular details of these processes, however, require further investigation to merge the observations with a mechanistic understanding.

Apart from experimental studies, theoretical and computational models have been used to study various aspects of X-inactivation. Early theoretical works used stochastic models to explore the initiation of XCI, which involves counting of X alleles, their pairing, and the choice of inactive X chromosome [41–45]. With the availability of chromatin conformation capture data (3C, 5C, and Hi-C), recent polymer modeling studies have used contact probability data to reconstruct 3D conformations of the X chromosome and *Xic* region on the Mb scale [9, 20, 46, 47]. These studies reveal the formation of megadomains during X-inactivation and how the chromosome organization changes during embryonic development. While these studies explore some aspects of XCI, a high-resolution mechanistic picture of how changes in gene interactions in the *Xic* of Xa and Xi regulate *Xist* expression is unavailable.

In this work, we address these knowledge gaps by studying how chromatin organization of inactive X differs from that of active X and how the interactions between key regulatory sequences and genes in the *Xic* region, including *Xist*, *Tsix*, *Jpx*, *Ftx*, *Rnf12*, *Linx*, and *Xite*, shape the inactivation process. While some of the sequences like *Jpx*, *Tsix*, and *Xite* are very close to the *Xist* gene (10s of kb), other genes like *Ftx*, *Linx*, and *Rnf12* are hundreds of kilobase-pairs apart.

To interpret the organization of *Xic* at these two different scales, we develop a multiscale modeling approach, where our validated nucleosome resolution mesoscale model is used to study the organization of a 90kb region consisting of *Xist*, *Tsix*, *Jpx*, and *Xite* [48, 49] (see Fig. 1, 90kb region highlighted in yellow). In parallel, we utilize a coarse-grained (CG) polymer model with an explicit loop extrusion mechanism to probe the organization of a 1 Mb region around *Xist*, as illustrated in Fig. 1. We incorporate epigenetic data from experiments that employ allele-specific design to distinguish the active X chromosome from the inactive X allele in our models (Fig. 2). Our nucleosome resolution model reveals key changes in the organization of the *Xite* enhancer, *Tsix* promoter (*Tsixp*), and *Xist* gene on Xa and Xi. We demonstrate how these changes correlate with their regulatory role in the XCI process. Our coarse-grained polymer model reveals that interactions between different genes and regulatory sequences change during X-inactivation. Specifically, we find that the loss of interactions between *Linx* and the *Xite*/*Tsixp*, as well as the formation of a transcriptionally active domain around *Xist*, *Jpx*, and *Ftx*, occurs after XCI. Together, integrating 3D structural insights from multiscale simulations with epigenetic features allows us to link the reorganization of the *Xic* region to its biological regulatory role in the XCI process and to propose some general principles of gene regulation.

**FIG. 2.**
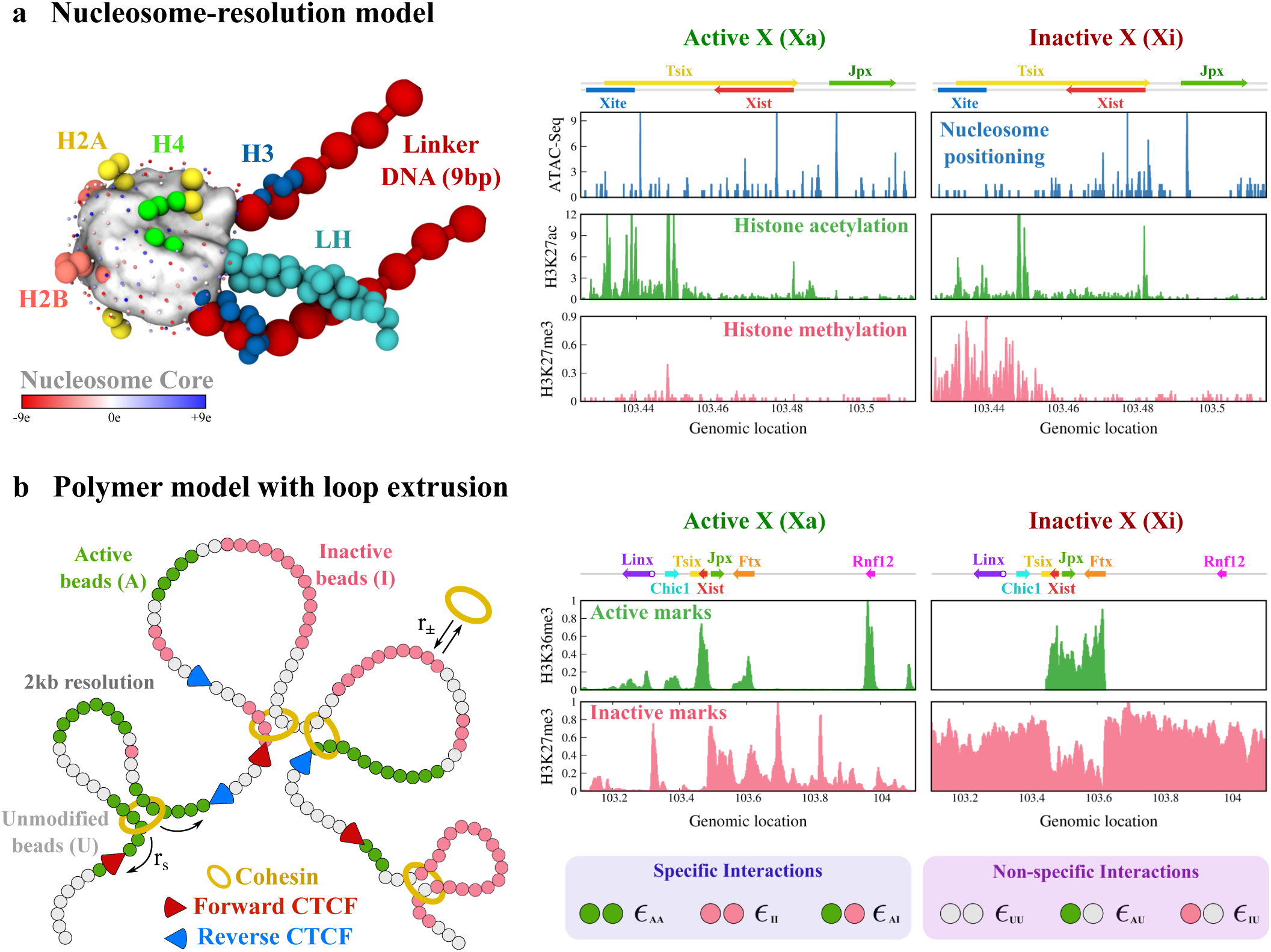
Nucleosome resolution model and polymer model with loop extrusion. **a** The nucleosome-resolution model consists of an electrically charged nucleosome core particle (300 discrete charges on the nucleosome surface), histone tails (H2A, H2B, H3, and H4), linker histone (LH), and linker DNA beads. We use ATAC-seq data from allele-specific experiments to assign nucleosome positions for Xa and Xi. Histone-modification data (acetylation and methylation) are incorporated as follows: H3K27ac data are used to model changes in the local folding of H3 tails, and H3K27me3 data are used to model long-range, protein-mediated contacts for Xa and Xi. **b** The polymer model with loop extrusion incorporates cohesin binding and dissociation (rate *r*_±_) and loop extrusion (rate *r_s_*) until cohesin is blocked by oppositely oriented, bound CTCF proteins or by another cohesin complex. For Xa and Xi, we use histone-modification data (H3K36me3 for active regions and H3K27me3 for inactive regions) to stochastically assign chromatin-bead types—active (A), inactive (I), or unmodified (U)—for each trajectory. Based on these modification marks, non-bonded chromatin-bead pairs are assigned different strengths of attractive interactions (*ɛ*).

## RESULTS

### Nucleosome-resolution model reveals epigenetically-driven locus-specific reorganization of *Xic* regulatory elements

To investigate local chromatin reorganization during X inactivation, we employed our nucleosome-resolution model to simulate a 90 kb region of *Xic*, consisting of key regulatory elements *Xite*, *Tsix*, *Xist*, and *Jpx* (Fig. 2a). This nucleosome resolution model has been validated against various experiments [49–57]. The model consists of a coarse-grained nucleosome core with point charges and beads for linker DNA, linker histones (LH), and histone tails (see Methods). The total energy consists of stretching, bending, and twisting terms for linker beads and the nucleosome core; stretching and bending terms for the histone tails and linker histone beads; excluded volume and electrostatic interactions among all particles (beads and nucleosome charges); and long-range interactions between methylated nucleosomes (Supplementary Note 1). We use equilibrium Monte-Carlo to sample chromatin conformations. Using allele-specific ATAC-seq data for Xa and Xi, we generate an ensemble of chromatin conformations, where each structure has unique nucleosome positions sampled from a life-like distribution of linker lengths (see Methods). Similarly, we use histone modification data (H3K27ac and H3K27me3) for Xa and Xi to model epigenetic interactions (Fig. 2a). Since allele-specific linker histone binding data were not available, we position LHs so as to keep the overall 50% occupancy in 90kb region (i.e., one LH per two nucleosomes) but increase occupancy to 100% at H3K27me3-marked regions, reflecting the established correlation between linker histone enrichment and repressive H3K27me3 marks [58].

Fig. 3a-b shows contact maps calculated from equilibrated nucleosome-resolution simulations of Xa and Xi, with regulatory sequences annotated on top. We observe a moderate increase in the contact probability around *Xite* locus (blue box) upon inactivation. Representative 3D conformations of the simulated chromatin fiber show an open conformation of *Xite* on Xa, while a more compact domain marked by H3K27me3 emerges for Xi (Fig. 3c-d and Supplementary Fig. 1-2).

**FIG. 3.**
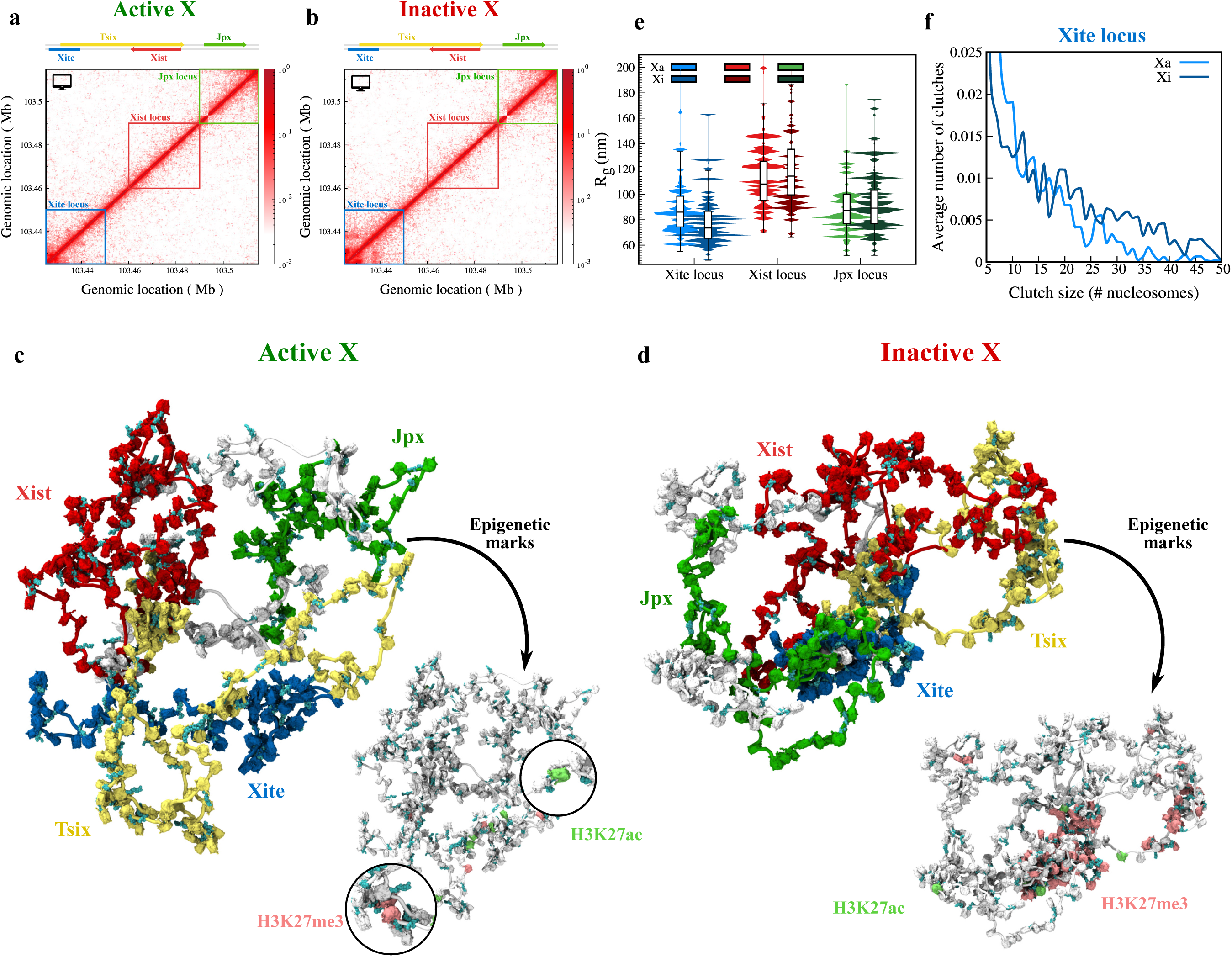
Nucleosome-level simulations reveal epigenetically driven compaction and clutch enlargement during X inactivation. **a-b** Internucleosome contact maps of the 90 kb region from nucleosome-resolution simulations are shown for (**a**) the active X and (**b**) the inactive X. Genes within this region are annotated at the top for reference. **c-d** Representative 3D conformations of folded *Xic* region are shown from converged MC simulations (each simulated for 200 copies and 10^8^ steps), for (**c**) Xa and **d** Xi. The same conformation is annotated by gene position (left) and by active and inactive epigenetic marks (right). **e** The radius of gyration of *Xite*, *Xist*, and *Jpx* loci (blue, red, and green boxes in **a-b**) is plotted for Xa and Xi as a violin plot. **f** Distribution of nucleosome clutch sizes in the *Xite* locus is shown for Xa and Xi. See Figs. 4 and 5 for details of Xite and Xist loci.

We used the radius of gyration (*R_g_*) to quantify global and locus-specific compaction. The *R_g_* of the entire 90 kb segment shows only a modest reduction on Xi compared to Xa (Supplementary Fig. 3). To probe local reorganization, we computed *R_g_* of individual *Xite*, *Xist*, and *Jpx* loci (blue, red, and green boxes in Fig. 3a-b). The *Xite* locus shows significant compaction upon inactivation (decreased *R_g_*), while *Xist* and *Jpx* show slight decompaction (increased *R_g_*) (Fig. 3e). This difference suggests that local organization is modulated by site-specific factors (epigenetic marks, nucleosome positioning, locus-bound regulatory complexes, etc.) rather than simple uniform compaction. This heterogeneity may reflect the diverse regulatory functions of these elements and their dynamic organization within *Xic*.

Further nucleosome clutch analysis in Supplementary Fig. 4a-b, reveals differences in nucleosome packing between Xa and Xi: Xa shows a higher number of small clutches, whereas Xi displays fewer but substantially larger clutches.

A clutch is a contiguous cluster of nucleosomes whose pairwise distance is below a threshold value (see Supplementary Note 2). Locus-specific clutch size distributions (Fig. 3f and Supplementary Fig. 4c-d) show a pronounced increase in large clutches at the *Xite* locus upon inactivation, while *Xist* and *Jpx* clutch distributions remain similar between Xa and Xi. This clutch enlargement at *Xite* reflects formation of a compact heterochromatic domain marked by H3K27me3 on Xi. The formation of these enlarged nucleosome groups arises primarily from increased nucleosome occupancy and recruitment of repressive complexes in methylated regions that promote nucleosome-nucleosome interactions.

### Nucleosome-resolution modeling reveals distinct reorganization of *Xite* and *Xist* loci during X-inactivation due to differences in epigenetic marks

To study locus-specific chromatin reorganization during XCI, we examined the structural changes at two loci: one encompassing the *Tsix* promoter (*Tsixp*) and its *Xite* enhancer, and another located around the *Xist* gene. Computed internucleosome 3D distance maps were compared with experimental ATAC-seq and histone modification profiles to link epigenetic landscapes and three-dimensional chromatin architecture to define mechanistic aspects (Figs. 4 and 5).

**FIG. 4.**
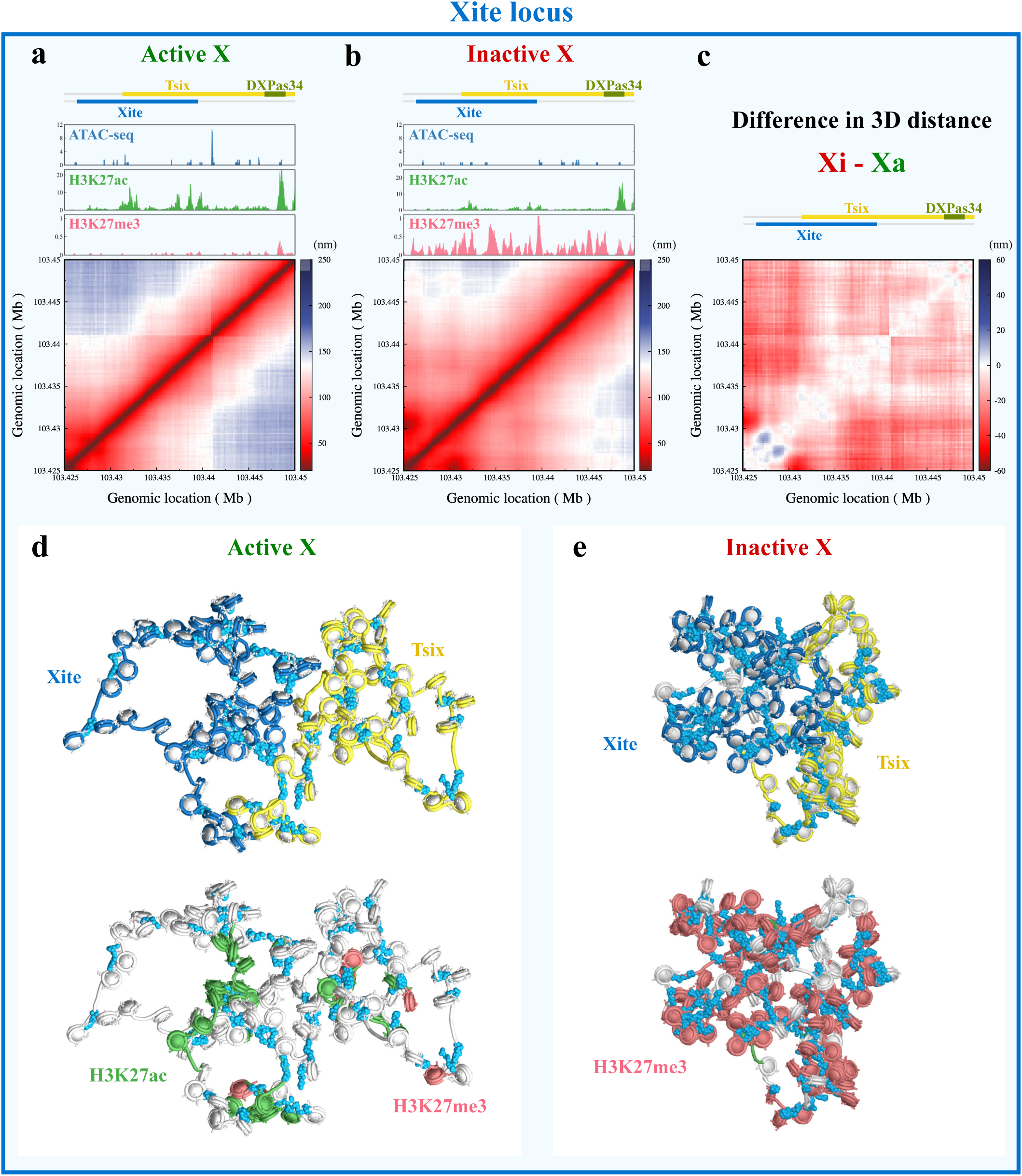
3D distance map analysis of nucleosome-resolution conformations reveals differential chromatin organization of *Xite* locus. **a-b** 3D distance maps from nucleosome-resolution model and corresponding experimental ChIP-seq profiles for the *Xite* locus are shown for (**a**) Xa and (**b**) Xi. ChIP-seq tracks include ATAC-seq (blue), H3K27ac (green), and H3K27me3 (red) data. **c** The difference in average 3D distance (Xi minus Xa) between nucleosomes is plotted as a heatmap with negative values shown in red and positive values shown in blue. **d-e** The representative 3D snapshots of the *Xite* locus are shown for (**d**) Xa and (**e**) Xi, with nucleosomes colored by gene position on the top and by histone marks on the bottom: H3K27ac (green) and H3K27me3 (light red). Turquoise units are linker histones (Fig. 2a).

**FIG. 5.**
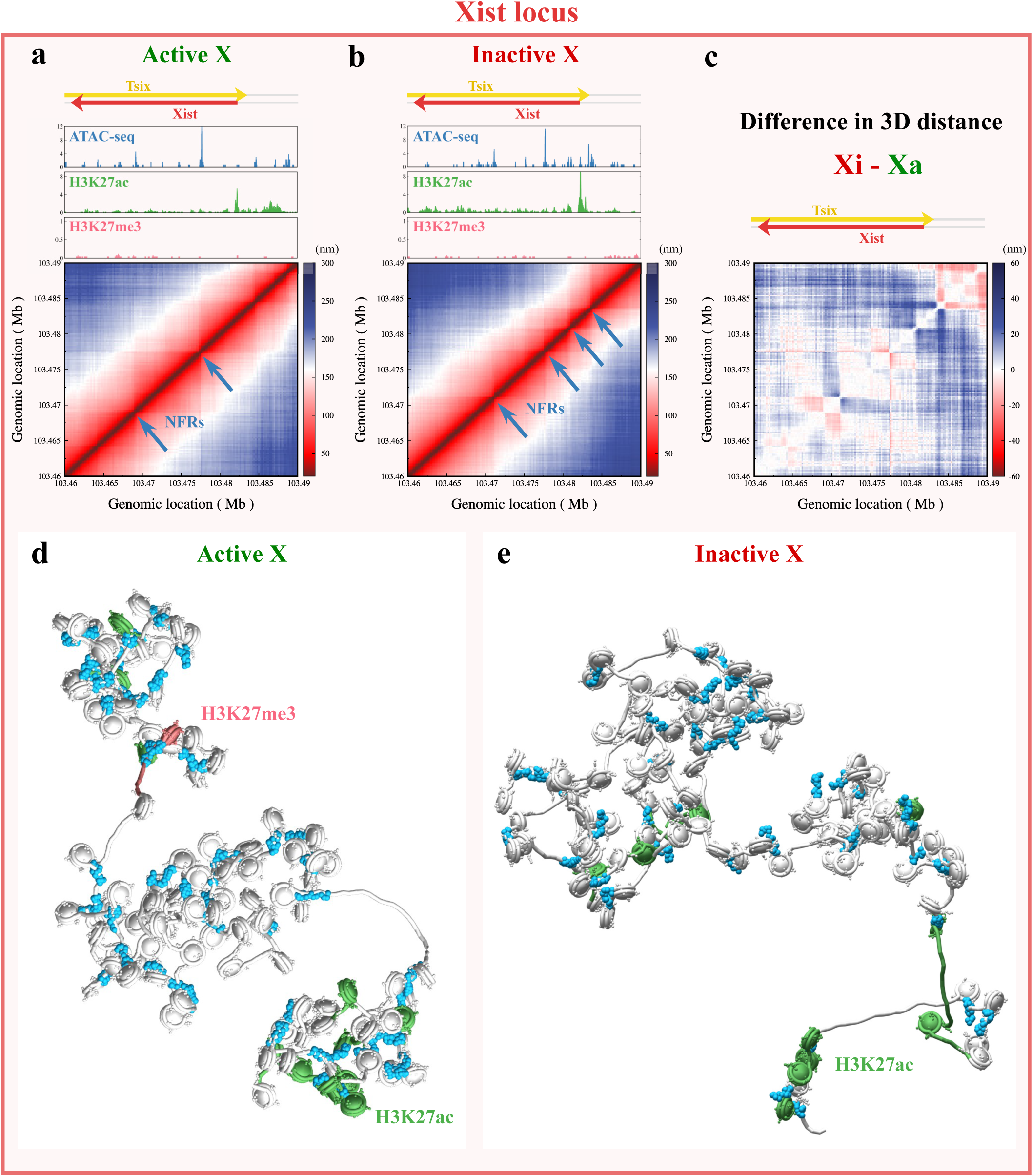
3D distance map analysis of nucleosome-resolution conformations reveals differential chromatin organization of *Xist* locus. **a-b** 3D distance maps from nucleosome-resolution model and ChIP-seq profiles for the *Xist* locus are shown for (**a**) Xa and (**b**) Xi. ChIP-seq tracks include ATAC-seq (blue), H3K27ac (green), and H3K27me3 (light red) data. The nucleosome-free regions (NFRs) from the ATAC-seq peak are marked on the contact maps using blue arrows. **c** The difference in average 3D distance (Xi minus Xa) between nucleosomes is plotted as a heatmap with negative values shown in red and positive values shown in blue. **d-e** The representative 3D snapshots of the *Xist* gene are shown for (**d**) Xa and (**e**) Xi, with nucleosomes colored by histone marks: H3K27ac (green) and H3K27me3 (light red). Turquoise units are linker histones (Fig. 2a).

The *Xite*/*Tsixp* region shows the most dramatic reorganization (Fig. 4a-b). On Xi, we observe the formation of a prominent heterochromatic domain, characterized by extensive enrichment of the repressive H3K27me3 mark relative to Xa. Fig. 4c shows the differential 3D distance map (Xi minus Xa), where red indicates compaction (decreased distances on Xi), blue indicates expansion (increased distances on Xi), and white indicates no change. This map reveals substantial compaction of the *Xite* locus, evident from the predominantly red coloring indicating shorter pairwise distances within this region on Xi compared to Xa. At the same time, histone acetylation—a hallmark of active chromatin—is substantially depleted around the *Xite* locus and around the adjacent DXPas34 microsatellite repeat, a known positive regulator of *Tsix* [59].

Fig. 4d-e shows the representative snapshots of the *Xite* locus for Xa and Xi, with acetylated nucleosomes colored green and methylated nucleosomes colored light red. This substantial decrease in acetylated nucleosomes and a significant increase in the number of contacts between methylated nucleosomes lead to a compact domain for Xi. This coordinated epigenetic remodeling creates a repressive chromatin environment that effectively silences *Tsix* on the Xi allele. Since *Tsix* functions as a negative regulator of *Xist*, its silencing is essential for stabilizing the inactive state. The formation of this domain likely involves the recruitment of Polycomb Repressive Complexes (PRC), which catalyze the spreading of H3K27me3 and the formation of a heterochromatic condensate that prevents *Tsix* reactivation and ensures stable *Xist* expression.

Next, the *Xist* locus shows subtle yet functionally significant architectural changes (Fig. 5a-b). On Xa, *Xist* is partitioned into relatively large domains separated by nucleosome-free regions, NFRs (Fig. 5 blue arrows). However, on Xi, *Xist* is organized into multiple smaller domains, separated by nucleosome-free regions, as illustrated by the experimental ATAC-seq curve at the top. This increased domain fragmentation may facilitate transcriptional activity by providing access for regulatory factors and RNA polymerase II. The differential 3D distance map (Fig. 5c, Xi minus Xa) reveals decompaction around the *Xist* gene, particularly near the *Xist* transcription start site (TSS), shown by the blue coloring indicating increased pairwise distances within this region on Xi compared to Xa. This more open structure at the *Xist* TSS, combined with increased levels of the active histone mark H3K27ac (Fig. 5d-e), likely promotes transcription factor binding to facilitate sustained transcription of *Xist*.

We also examined the *Jpx* gene, whose non-coding RNA is known to positively regulate *Xist* expression in Supplementary Fig. 5. The minimal changes in chromatin organization for the *Jpx* locus between Xa and Xi could be due to partial expression of *Jpx* on Xa with higher expression levels on Xi [22]. Unlike *Tsix*, which must be completely silenced for XCI to proceed, *Jpx* maintains a permissive chromatin state on both chromosomes. This differential reorganization—where *Xite*/*Tsixp* undergoes heterochromatin compaction, *Xist* adopts a more open, fragmented structure, and *Jpx* remains largely unchanged—highlights the selective nature of chromatin reorganization during XCI.

### Polymer modeling reveals how loop extrusion and epigenetic interactions reorganize *Xic* during inactivation

To probe chromatin reorganization at a broader genomic scale, we developed a coarse-grained polymer model (2 kb resolution) of a 1 Mb region surrounding *Xist*. This extended region includes several genes and regulatory elements such as *Linx*, *Chic1*, *Ftx*, and *Rnf12*, as shown in Fig. 1. The polymer model with loop extrusion enables us to study long-range regulatory interactions beyond the central 90kb region. Our model integrates two fundamental mechanisms of genome organization: CTCF/cohesin-mediated loop extrusion and phase separation driven by epigenetic interactions (Fig. 2b). Polymer beads were designated as active or inactive based on H3K36me3 and H3K27me3 histone modification profiles, respectively. Polymer beads with identical marks were assigned attractive self-interactions, recapitulating the phase-separation behavior observed for chromatin domains with similar epigenetic states. Loop extrusion was implemented using a stochastic model, where cohesin complexes bind and dissociate at rates *r*_±_ and extrude the loop bidirectionally at a speed *r_s_* until encountering convergent CTCF sites or colliding with other extruding cohesins [60]. CTCF binding probabilities across individual simulation trajectories were drawn from experimental ChIP-seq profiles, capturing the stochastic nature of CTCF occupancy and its allele-specific binding patterns (see Methods).

Fig. 6a calibrates the strength of epigenetic interactions to ensure our model faithfully captures the features from the high-resolution Micro-C contact map of the active X in mESCs. The simulated contact map (Fig. 6a bottom) correlates strongly with the experimental map on top (stratum-adjusted correlation = 0.83; Fig. 6a and SI Supplementary Fig. 6). These optimized parameters, determined only from Xa data, were then used without further adjustment to simulate Xi chromatin organization using allele-specific CTCF ChIP-seq and histone modification data from female MEFs.

**FIG. 6.**
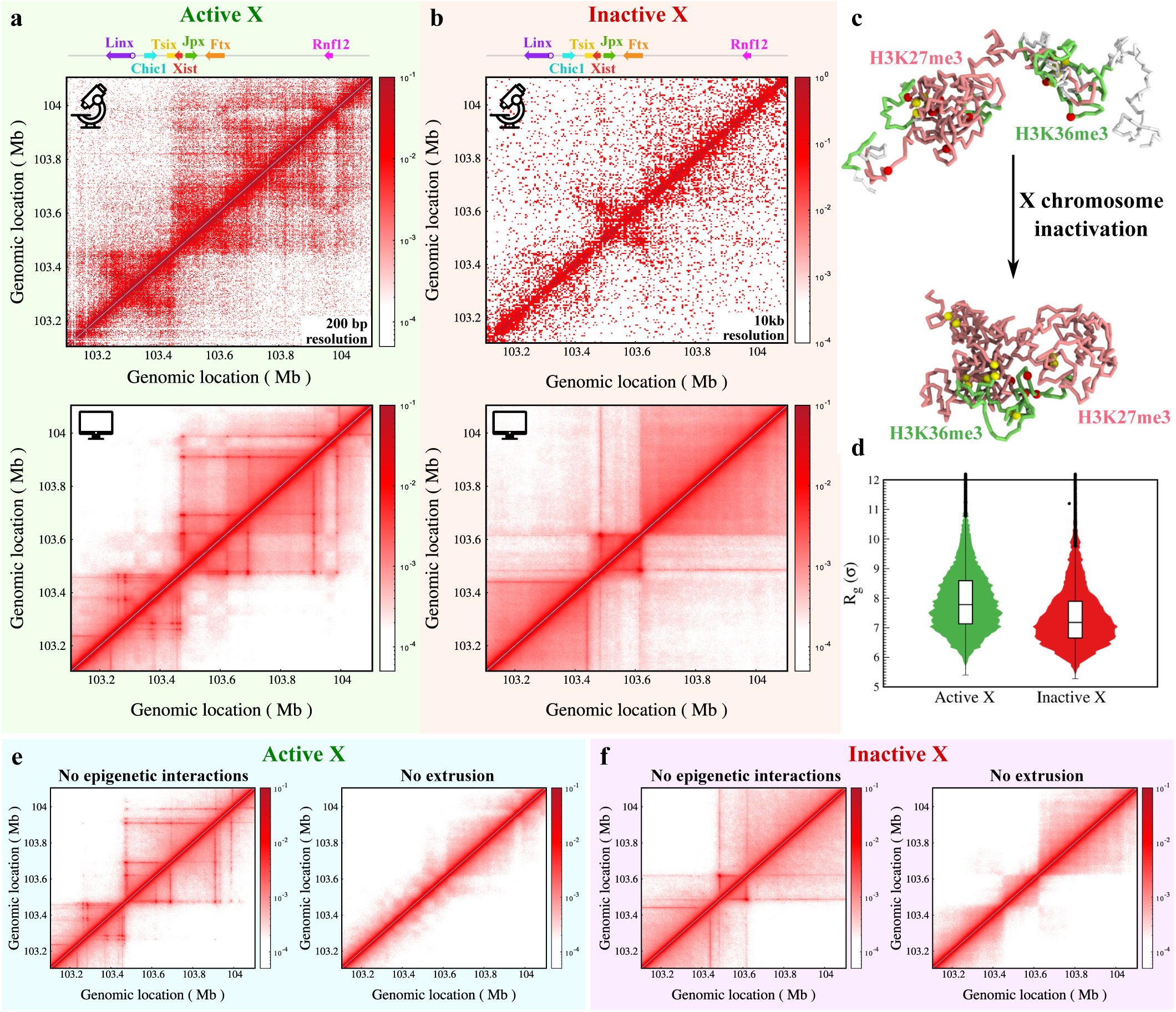
Loop extrusion and epigenetic interactions together contribute to reorganization of *Xic* during XCI. **a–b** Experimental contact maps from Hsieh et. al. [61] and Du et. al. [34] (top row with microscope symbol) of the 1 Mb region are compared with simulation contact maps generated using the polymer model (bottom row with computer symbol) for (**a**) the active X and (**b**) the inactive X. The genes located in this region are annotated on top for reference. **c** The representative snapshots of the simulated region are shown for Xa and Xi, with H3K36me3 marked regions shown in green and H3K27me3 marked regions shown in light red. **d** The overall radius of gyration of the 1Mb region is plotted for Xa and Xi as a violin plot. **e-f** The contact map of the *Xic* is plotted in the absence of epigenetic interactions (left) and in the absence of loop extrusion (right) for (**e**) Xa and (**f**) Xi.

Despite the lower resolution of available experimental Xi contact maps (10 kb bins), our simulations capture key architectural features observed experimentally (Fig. 6b). The successful prediction of Xi architecture using Xa-derived parameters suggests that XCI-associated structural reorganization primarily arises from altered CTCF binding patterns and epigenetic landscapes, rather than changes in fundamental biophysical interactions.

Fig. 6c shows representative polymer snapshots from steady-state conformations, with active H3K36me3-marked regions shown in green and inactive H3K27me3-marked regions shown in light red. We observe self-segregation of active (green) and inactive (light red) chromatin beads, with cohesin anchor sites (yellow) and CTCF boundaries (red) forming architectural hubs. The polymer also exhibits a compact overall structure after inactivation. We quantify the compaction by computing the *R_g_* of the whole 1MB region for Xa and Xi (Fig. 6d). Consistent with nucleosome-resolution modeling, the Xi model showed a lower overall *R_g_* than Xa, confirming hierarchical compaction across scales—from nucleosome clutches to megabase-level folding.

To demonstrate that both loop extrusion and epigenetic marks are necessary to capture the experimental trend for *Xic*, we remove epigenetic interactions and loop extrusion in turn. With only loop extrusion (no epigenetic interactions), the overall TADs and large-scale loop structures on both Xa and Xi are retained (Fig. 6e-f). However, the finer-scale contact enrichments within TADs on Xa and inter-TAD interactions on Xi cannot be captured; these features emerge from compartmentalization driven by epigenetic interactions. Conversely, with epigenetic interactions and no loop extrusion, diffused contact patterns lacking defined TAD structures and CTCF-anchored loops emerge. Thus, loop extrusion establishes the structural scaffold of TAD organization, while epigenetic interactions modulate intra- and inter-TAD contact frequencies through the process of compartmentalization. Importantly, neither mechanism is sufficient to reproduce experimental contact maps on its own; instead, the *Xic* architecture emerges from their interplay. This interplay may represent a general organizing principle for mammalian chromatin, where extrusion forms long-range loops while epigenetic interactions form regulatory hubs and compartments.

### Regulatory interactions are differentially rewired between active and inactive X chromosomes

Next, we use the 3D chromatin conformations generated by our coarse-grained polymer model to investigate how regulatory contacts within the *Xic* are rewired during XCI. In Fig. 7a-b, we plot the contact probability of specific genes and regulatory sequences (*Xite*, *Tsix*, *Xist*, and *Jpx*) with all other genomic regions in the *Xic* to highlight changes in their contact interactions from Xa to Xi. The genes and regulatory sequences in the region are annotated at the top. The contact interactions for Xa from our simulations in Fig. 7a compare well with experimental observations, validating our model. Meanwhile, the predictions for Xi enable us to investigate changes in regulatory interactions between different genes.

**FIG. 7.**
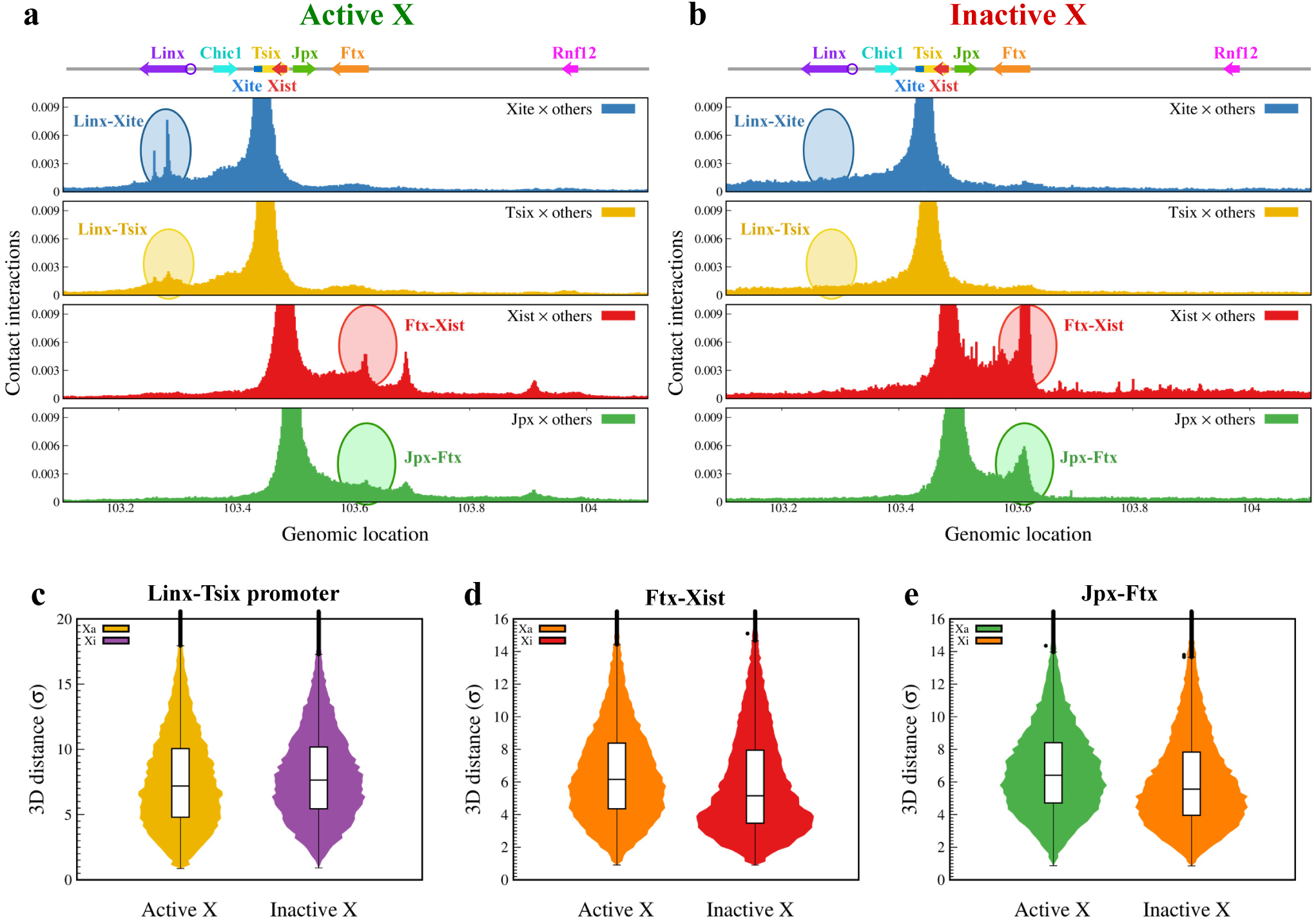
Contact interactions and 3D distances reveal rewiring of regulatory networks in *Xic* during XCI **a-b** Interaction profiles of transcription start sites (TSSs) of key genes and regulatory elements with other genomic locations are shown for (**a**) Xa and (**b**) Xi. Gene annotations are displayed along the top, and major interaction changes between the active and inactive X are highlighted with colored ovals. **c-e** Pairwise 3D distances between regulatory loci—(**c**) *Linx* -*Tsixp*, (**d**) *Ftx* -*Xist*, and (**e**) *Jpx* -*Ftx* —highlight spatial reorganization during X inactivation.

On Xa, the *Linx* locus shows strong spatial contacts with the *Tsix* promoter and the *Xite* enhancer, while on Xi, these interactions are completely lost (Figs. 7a-b *Xite* and *Tsix* panels). Experiments have suggested that the *Linx* gene acts as a positive regulator of *Tsix*, and our results provide a structural mechanism for this regulation: *Linx* may activate the *Tsix* gene *in cis* by spreading the lncRNA through spatial proximity on Xa. After X-inactivation, these interactions are lost, with *Linx* becoming spatially segregated from the *Tsix* /*Xite* regulatory hub. The corresponding increase in 3D distance between *Linx* and the *Tsix* promoter (Fig. 7d) confirms that the physical decoupling of these regulatory sequences coincides with *Tsix* silencing. This effectively isolates the *Tsix* /*Xite* hub from its upstream activators, likely contributing to the robust silencing of *Tsix* on the Xi.

The most prominent architectural reorganization in *Xic* involves the formation of an active chromatin domain marked by H3K36me3 containing the loci *Xist*, *Jpx*, and *Ftx* on Xi (Fig. 7b *Xist* and *Jpx* panels). This active compartment formation significantly enhances spatial interactions among these three genes, resulting in a substantial increase in *Xist* -*Ftx* contact frequency (*Xist* panel in Fig. 7a-b). The *Ftx* -*Xist* distance distribution on Xi is strongly skewed toward smaller values as compared to Xa (Fig. 7c). Similarly, *Ftx* and *Jpx* —both known positive regulators of *Xist* —exhibit reduced spatial separation on Xi relative to Xa, reflecting their co-localization within the active compartment. This active domain may represent a dedicated transcriptional factory or enhancer hub that maintains the exceptionally high levels of *Xist* expression necessary for chromosome-wide silencing. By concentrating *Xist*, *Jpx*, and *Ftx* in a confined nuclear territory, this compartment may facilitate quick access to regulatory factors and coordinate the expression of multiple pro-inactivation lncRNAs to stabilize *Xist* transcription.

## DISCUSSION

Our multiscale computational approach provides a biophysical framework for the complex architectural and regulatory rewiring of the *Xic* during X inactivation. We have integrated the findings from nucleosome-resolution simulations and coarse-grained polymer modeling to uncover how the interplay between loop extrusion and epigenetic modifications modulates differential regulatory patterns in Xa and Xi. While many experiments have predicted the regulatory roles of different genes in the *Xic*, our models provide a mechanistic view of how epigenetic interactions and loop extrusion rewire these interactions during XCI. Moreover, our approach provides high-resolution chromatin conformations of *Xic* as compared to previous modeling approaches [20, 47].

Both nucleosome-resolution and polymer simulations show an increase in compaction of *Xic* following X-inactivation. The compaction of the 90kb region is accompanied by a shift in nucleosome packing with the transition from numerous small nucleosome clutches on Xa to fewer, substantially larger clutches on Xi (Fig. 3 and Supplementary Fig. 4). The increase in nucleosome clutch size is predominantly around the *Xite* locus, likely due to nucleosome bridging through PRC complexes and increased nucleosome occupancy [62, 63]. This structural change is triggered by the local epigenetic landscape, with the most significant changes observed around the *Xite*/*Tsixp* region, where a heterochromatic domain is formed (Fig. 4). The formation of this domain on Xi is characterized by an increase in H3K27me3 marks and the loss of activating H3K27ac marks (Fig. 8). Our coarse-grained polymer simulations on the 1Mb scale extend this picture by revealing that the *Xic* undergoes long-range spatial reorganization, losing its bipartite TAD structure (Fig. 8). This results in the *Xite*/*Tsixp* losing interactions with the upstream activator *Linx*. Namely, the strong contacts of *Linx* with *Tsix* on Xa are lost upon inactivation (Fig. 7), explaining *Linx* ’s known role as a positive regulator of *Tsix* [18, 20, 64]. The combination of local heterochromatin formation and long-range spatial segregation may provide robust silencing mechanisms that ensure stable *Tsix* repression essential for sustained *Xist* expression on Xi.

**FIG. 8.**
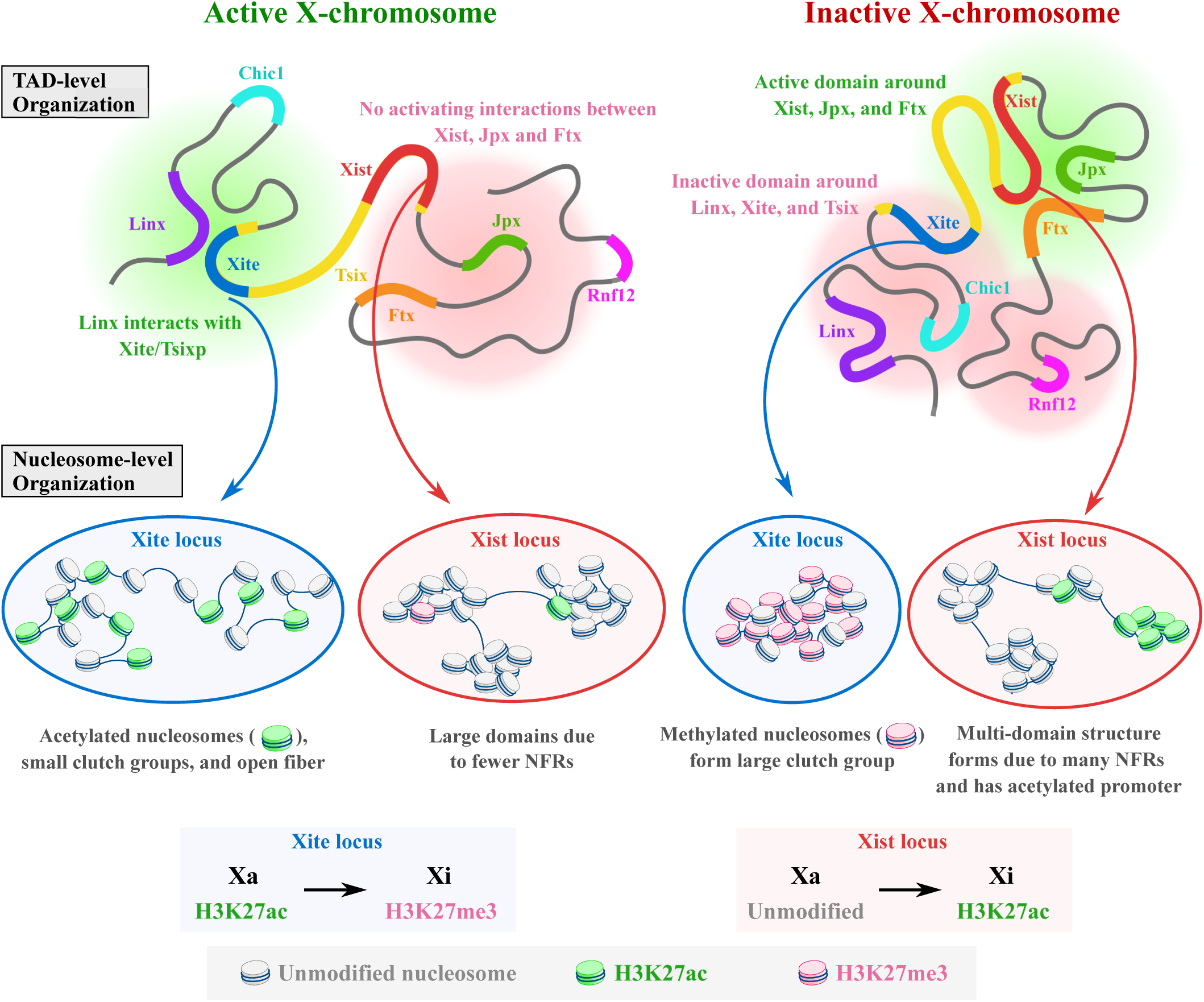
Schematic summary of multiscale modeling insights into *Xic* reorganization during XCI. At the scale of TADs, Xa exhibits strong *Linx* –*Xite*/*Tsixp* interactions that are essential for *Tsix* expression. These contacts are lost upon inactiva-tion, and *Linx*, *Xite*, and *Tsix* become incorporated into an inactive domain (red cloud). In contrast, the *Xist*, *Jpx*, and *Ftx* loci—which show no activating interactions Xa—coalesce into a unified active compartment on Xi (green cloud), promoting enhanced contacts among these known Xist activators. These large-scale changes are accompanied by finer nucleosome-level reorganization captured by our nucleosome-resolution modeling. The *Xite* locus, initially marked by active H3K27ac modifi-cations and smaller clutches, compacts into a heterochromatic domain (H3K27me3 marks) with enlarged clutches consistent with transcriptional silencing. Meanwhile, the *Xist* locus transitions from a structure with large domains on Xa to an open multi-domain architecture with more H3K27ac-marked nucleosomes near the *Xist* promoter region on Xi.

While *Tsix* undergoes comprehensive silencing through the formation of a heterochromatic domain and spatial isolation from *Linx*, the *Xist* locus shows a fragmented multi-domain architecture on Xi, characterized by multiple smaller domains separated by extended nucleosome-free regions (Figs. 5 and 8). We observe decompaction of the region proximal to the *Xist* TSS on Xi with elevated levels of histone acetylation. This active domain likely promotes recruitment of transcriptional machinery to sustain the high expression levels of *Xist*. Unlike *Xite* and *Xist*, the *Jpx* locus shows a similar organization for Xa and Xi, likely due to partial expression of *Jpx* on Xa [22] and the need for an additional mechanism, such as loop extrusion, that could facilitate *Jpx* -*Xist* interaction. Our polymer simulations with loop extrusion fill this gap by showing how *Xist* becomes a part of an H3K36me3-enriched active compartment encompassing *Jpx* and *Ftx*, thereby spatially organizing *Xist* with its known transcriptional activators [22, 24] (Fig. 8). The formation of this active domain increases the spatial interactions between these genes on Xi, with a decrease in *Xist* -*Ftx* and *Jpx* -*Ftx* distances (Fig. 7d-e). This differential organization of genes on the nucleosome level and rewiring of regulatory interactions on the TAD level highlight hierarchical multiscale reorganization of chromatin architecture during XCI.

Another mechanistic insight from our polymer simulations is that both CTCF-cohesin-mediated loop extrusion and epigenetic interactions work together to drive reorganization of *Xic* locus on Xa and Xi. This finding is consistent with the observations from cohesin depletion experiments, which eliminate TADs but preserve compartmentalization [65, 66]. In the context of XCI, loop extrusion establishes the structural scaffold that constrains where epigenetic domains can form, while compartmentalization—driven by self-attraction of similarly modified chromatin—modulates contact frequencies within and between TADs. The specific changes in CTCF binding during XCI thus reorganize TAD structure by tuning loop positions to facilitate formation of the *Tsix* -*Xite*-*Linx* heterochromatic domain and the *Xist* -*Jpx* -*Ftx* active domain. These resulting active and inactive domains resemble phase-separated chromatin condensates, similar to transcriptional condensates near super-enhancers and PRC-driven heterochromatic droplets, with loop extrusion acting to stabilize and maintain their structural boundaries. Importantly, we demonstrate that the parameters optimized from Xa data successfully predict the Xi architecture, indicating that the fundamental biophysical principles governing chromatin folding are robust.

Beyond advancing our understanding of XCI, our findings shed light on general principles of gene regulation through chromatin reorganization. The spatial insulation mechanism of regulatory elements, segregated from their target genes to enforce silencing, may explain how cells prevent inappropriate enhancer-promoter interactions during differentiation without requiring complete enhancer inactivation. On the other hand, the clustering of positive regulators into a shared transcriptional domain is similar to the organization of super-enhancers, suggesting a general mechanism for achieving robust, sustained gene expression through phase separation and spatial proximity. The hierarchical nature of XCI-associated reorganization, spanning nucleosome clutches to megabase-scale compartments, suggests that cells employ multiple mechanisms that actively maintain specific three-dimensional structures that reinforce transcriptional programs and create epigenetic memory.

## METHODS

We employ a multiscale modeling approach to investigate chromatin reorganization within the *Xic* during X-chromosome inactivation. Our strategy integrates two complementary models operating at different length scales (Fig. 1). Our nucleosome-resolution model explicitly incorporates individual nucleosome cores, histone tails, linker DNA, and linker histone proteins to simulate a 90 kb region consisting of regulatory elements *Xist*, *Xite*, *Tsix*, and *Jpx* (Fig. 2a). This high-resolution approach captures the changes in local chromatin folding of these critical loci during XCI. Our coarse-grained polymer model with loop extrusion simulates the larger 1 Mb *Xic* region, including additional regulatory genes *Linx*, *Chic1*, *Ftx*, and *Rnf12*, enabling analysis of long-range chromosomal interactions and domain organization (Fig. 2b).

### Nucleosome resolution model

We utilize our previously validated model to simulate chromatin at nucleosome resolution [49–57]. This model comprises four main components: nucleosome core, linker DNA, histone tails, and linker histone proteins. The former is represented by point charges to mimic the electrostatic environment of the nucleosome without the histone tails. The others (linker DNA, histone tails, and linker histone) are denoted by coarse-grained (CG) beads. The model also incorporates the biophysical effects of histone tail modifications (H3K27ac and H3K27me3). Each of these components is modeled with a specific modeling strategy, which is then integrated together to simulate the *in vivo* chromatin polymer. Below, we provide a brief description of the modeling approach for each of these components, with further details provided in earlier works.

#### Nucleosome core particle

Each canonical nucleosome core particle in our model consists of a histone octamer (H2A, H2B, H3, and H4) and ∼ 146bp of DNA wrapped around it. We use *N_c_* = 300 discrete charged particles uniformly distributed on an irregular rigid surface to simulate the nucleosome core. The charges on these particles are optimized using the Discrete Surface Charge Optimization (DiSCO) algorithm such that the electric field, calculated using the Debye-Hückel approximation, from these charges mimics the field from atomistic simulations of the nucleosome core [50]. Because the nucleosome core is a relatively rigid structure, we use a rigid-body representation to model its dynamics.

#### Linker DNA

We use a bead-spring representation with stretching, bending, and twisting energy terms to model the linker DNA [67]. Each linker DNA bead represents ∼ 9bp DNA with a 3nm length. The negative charge on these linker DNA beads is assigned according to the physiological salt concentration. A 5bp DNA bead model was also introduced recently [68].

#### Histone tails and histone modifications

While the nucleosome core particle includes the structured part of the histone octamer core, the intrinsically disordered histone tails are modeled as a flexible bead-spring polymer. Each CG bead represents 5 amino acid residues with N-terminal tails of H2A, H2B, H3, and H4 having 3, 5, 8, and 5 CG beads, respectively, and 3 beads for the C-terminal tail of H2A [69]. We also model the effects of histone acetylation (H3K27ac) and histone methylation (H3K27me3) on these tails. Histone acetylation is modeled as folded tails with enhanced rigidity, as validated by previous studies [52, 53]. For the histone methylation mark, we use a machine learning algorithm to predict the contacts between nucleosomes with histone methylation marks [70]. The machine learning algorithm was trained on experimental data with and without methylation marks to predict methylation-mediated internucleosomal interactions.

#### Linker histone

The flexible linker histone is modeled at the same resolution as histone tails, with the globular head (GH) consisting of 6 beads and the C-terminal domain (CTD) having 22 beads [51]. Linker histones interact with linker DNA and stabilize compact chromatin structures through electrostatic interactions.

### Simulation Protocol

Based on the four components of our nucleosome-resolution model, we define the total potential energy of the system as

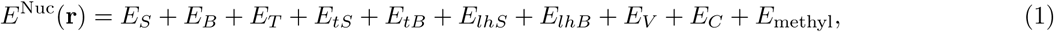

where *E_S_*, *E_B_*, and *E_T_* denote the stretching, bending, and twisting energies of the linker DNA beads and nucleosome core. The terms *E_tS_* and *E_tB_* describe the stretching and bending contributions of the histone tails, while *E_lhS_* and *E_lhB_* represent the corresponding stretching and bending energies of the linker-histone beads. Short-range excluded-volume interactions between all particles are captured by *E_V_*, and electrostatic interactions among charged particles are included in *E_C_*. The final term, *E*_methyl_, accounts for long-range contacts between H3K27me3-modified nucleosomes predicted by a machine-learning model trained on the ChIP-seq and Hi-C data (with and without methylation) [70]. We sample equilibrium chromatin configurations using a Metropolis Monte Carlo (MC) algorithm that incorporates both local and global trial moves. Local moves involve translation and rotation of a randomly selected particle. Global moves are implemented by selecting a random particle and rotating the shorter segment of the chromatin fiber around a randomly oriented axis passing through that particle, with the rotation angle drawn from a uniform distribution. The magnitude of the local displacement and the bounds of the global rotation angle are tuned to maintain an acceptance ratio of approximately 50%. After each proposed move, we calculate the associated change in energy and accept or reject the move based on the Metropolis criterion [71]. We also perform the Tail regrowth move, which involves regrowing a randomly chosen histone tail bead by bead using the Rosenbluth scheme.

Simulation trajectories were initialized from random, self-avoiding configurations in which straight linker DNA segments were assigned and nucleosome cores were placed with random orientations constrained by a 90^◦^ entry–exit angle. For both the active and inactive X chromosomes, we generated 200 independent trajectories. Each trajectory was equilibrated for 10^8^ Monte Carlo steps (MCS), and equilibration was confirmed by monitoring the time-dependent convergence of the radius of gyration (*R_g_*). After equilibration, we sampled 100 chromatin conformations from each trajectory at intervals of 50,000 steps to compute average quantities and distributions.

#### Modeling of the 90 kb *Xic* region for Xa and Xi using epigenomic data

To construct nucleosome-resolution models of active (Xa) and inactive (Xi) X chromosomes, we integrate multiple allele-specific epigenomic datasets that can distinguish Xa from Xi. Generally, this requires: (1) nucleosome positioning from ATAC-seq, (2) H3K27ac ChIP-seq profiles, (3) H3K27me3 ChIP-seq profiles, and (4) linker histone binding data. Such comprehensive allele-specific datasets are rare. We therefore integrate data from multiple experimental sources to model Xa and Xi. We primarily use allele-specific ATAC-seq and H3K27me3 profiles generated in neural progenitor cells (NPCs). However, allele-specific H3K27ac data are not available for NPCs. To approximate the active histone landscape, we incorporate H3K27ac profiles from the TX1072 X-inactivation study, assuming that the establishment of acetylation marks after XCI follows similar principles across these related cell types.

For Xa, nucleosome positions are assigned using a life-like distribution of nucleosome repeat lengths (NRL) derived from genome-wide measurements [72], with extended nucleosome-free regions (NFRs) placed at ATAC-seq accessibility peaks. For Xi, we modify this NRL distribution to favor shorter linker lengths, reflecting the experimentally observed increase in nucleosome density on the inactive X chromosome [62, 63], while maintaining nucleosome-free regions (NFRs) identified by ATAC-seq. Histone acetylation is assigned to nucleosomes where the H3K27ac ChIP-seq signal exceeds a threshold of 1.5 over the background noise. Methylation-mediated internucleosomal contacts are predicted using our validated machine learning algorithm [70], which takes H3K27me3 ChIP-seq data as input. We normalize the combined methylation data for the 90kb region from Xa and Xi, and we constrain each nucleosome to form at most one methylation-mediated contact, reflecting the multivalent nature of polycomb-mediated bridging of nucleosomes. Since the allele-specific LH binding data were not available, we assigned the LH positions to maintain an overall 50% occupancy for the 90kb region, but increase the LH occupancy at H3K27me3-enriched regions to 100% based on the established positive correlation between linker histone binding and repressive chromatin marks [58].

### Coarse-grained polymer model with loop extrusion

To simulate the 1Mb region of the *Xic*, we use a bead-spring polymer representation for chromatin with *N* = 500 CG beads (2kb resolution) connected by harmonic springs. We treat chromatin as a flexible polymer at this resolution [73, 74]. Additionally, we consider the loop extrusion activity and intra-chromatin interactions between regions with active and inactive histone marks [60]. The total energy of the polymer is given by

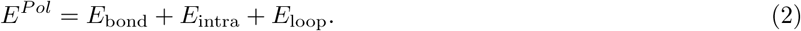

Here, the first term *E*_bond_ represents the bonding energy between consecutive beads modeled through a harmonic potential with equilibrium bond length *r*_0_ and spring constant *k_s_*. The second term denotes the non-bonded intra-chromatin interactions. We use the Lennard-Jones (LJ) potential to model these interactions. The strength of these interactions depends on the modification state of the corresponding beads. The strength of the self-interactions is higher compared to the interactions between beads having different modifications. The last term models the loop contact between the two anchor sites of cohesin complexes.

#### Epigenetic interactions

Our model has three types of epigenetic states for chromatin beads, active (A), inactive (I), and unmodified (U). For each trajectory, we assign the modification state to each chromatin bead stochastically, depending on the histone modification ChIP-seq profiles for H3K36me3 (active) and H3K27me3 (inactive) marks from experiments [34, 39]. We chose these modifications to assign active and inactive marks because of the higher correlation of the histone modification profile with the Hi-C pattern observed in experiments. We model these interactions using the truncated LJ potential

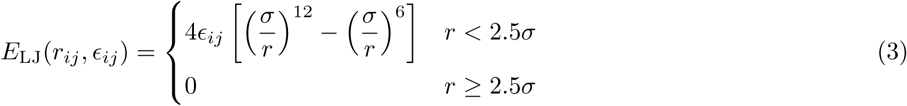

Here, *σ* is the diameter of the chromatin bead and *ɛ_ij_* represents the strength of attractive interactions between beads *i* and *j* depending on their type.

#### Loop extrusion activity

We also model the loop extrusion mechanism using kinetic events of binding/dissociation (*r*_±_) of cohesin complexes and the sliding (*r_s_*) of two anchor sites of cohesins in opposite direction (see Supplementary Note 1). The sliding of bound cohesins continues until they are blocked by oppositely oriented bound CTCF proteins or by another cohesin. We take the positions of CTCF binding locations from experimental ChIP-seq data. These CTCF sites are occupied by the CTCF proteins probabilistically in each trajectory, depending on their binding strength. The binding/dissociation rate of cohesin in our model allows us to control the mean number of cohesins bound (*N_c_*^0^). The loop formed by the sliding of the two anchor sites of each cohesin is modeled using a harmonic spring (last term in Eq. 2) with an equilibrium bond length *r*_0_ = *σ* and a spring constant *k_s_*.

#### Simulation protocol

We use the Metropolis Monte Carlo algorithm to simulate the coarse-grained polymer model with loop extrusion. Each Monte Carlo step (MCS) involves *N* displacement moves, where a random displacement vector is added to a randomly chosen bead. Then we compute the change in energy of the system using Eq. 2 and accept or reject the move based on the Metropolis criterion [71]. Additionally, we incorporate kinetic events of binding and dissociation, as well as the sliding of cohesin complexes, with rates *r*_±_ and *r_s_*, as described above. The positions of cohesin anchor sites are changed stochastically based on these kinetic rates. We simulate 120 trajectories for each parameter, equilibrate the system for 10^6^ steps, and sample the configurations every 5,000 steps for 5 × 10^6^ steps.

#### Modeling of the 1 Mb *Xic* region for Xa and Xi

Our coarse-grained polymer model assigns active and inactive bead types based on H3K36me3 and H3K27me3 ChIP-seq profiles, while CTCF binding probabilities are derived from CTCF ChIP-seq data. For Xa, we optimize interaction strengths using high-resolution Micro-C contact maps from mouse embryonic stem cells (mESCs)[61], achieving quantitative agreement between simulated and experimental contact maps. For Xi, we use these optimized parameters and using allele-specific H3K27me3 and CTCF ChIP-seq data from female mouse embryonic fibroblasts (MEFs). Since, allele-specific H3K36me3 data for MEFs are unavailable, we assign enriched H3K36me3 peaks around *Xist*, *Jpx*, and *Ftx* genes (Fig. 2b), based on the observed H3K36me3 profiles for Xi in extraembryonic ectoderm (ExE) cells [34].

#### Parameters

Polymer simulations were performed in dimensionless units, where lengths are expressed in bead diameters (*σ*) and energies in units of *k_B_T* . The polymer consists of *N* = 500 beads. Adjacent monomers, as well as pairs of cohesin anchor sites, are connected by harmonic springs with spring constant *k_s_* = 100 and equilibrium bond length *r*_0_ = *σ*. The simulation time was mapped to real time (1 MCS = 1 ms) by comparing the relaxation time of the polymer with the experimentally measured relaxation time of chromatin polymer [60]. The corresponding rate values used in real units are *r_s_* = 200bp*/*s and *r*_±_ = 5 × 10^−3^ s^−1^. The strength of attractive interactions between different types of chromatin beads was set to *ɛ_AA_* = *ɛ_II_* = 0.5, *ɛ_AI_* = 2, and *ɛ_UX_* = 0.3. Here, X refers to any other bead type (A, I, or U).

## DATA AVAILABILITY

Published datasets used in this study were obtained from the Gene Expression Omnibus (GEO) and ENCODE databases under the following accession codes: GSM1828646 (ATAC-seq), GSM3267061 (H3K27ac), GSM2667232/ ENCFF300QWW/GSM3040189 (H3K27me3 for NPC, mESC, and MEF cells, respectively), ENCFF300QWW (H3K36me3), GSE98671/GSE206011 (CTCF ChIP-seq for mESC and MEF cells, respectively), GSE130275 (Micro-C for mESC), and GSM6238884 (Hi-C for MEF). Relevant data generated from this study are included in this article’s Figures, text, and supplementary information.

## ACKNOWLEDGEMENTS

This work was supported in large part by the National Science Foundation Award 2337391 to T.S. Additional support was provided by the National Institutes of Health, National Institutes of General Medical Sciences Award R35-GM122562, the National Science Foundation Awards 2151777 and 2330628 from the Division of Mathematical Sciences and Award 2337391 from the Molecular Cellular Biosciences Core Programs, and Philip-Morris International to T.S. Support from the Simons Center of Computational Physical Chemistry at NYU to T.S. is also gratefully acknowledged (Award No. MPS-T-MPS-00839534 to Mark E. Tuckerman). Computing was performed on the Greene and Torch HPC clusters at New York University. We thank the NYU IT for providing resources, services, and expert advice. S.K. thanks Stephanie Portillo and Zilong Li for helpful discussions.

## AUTHOR CONTRIBUTIONS

S.K. designed the research, performed Monte Carlo simulations and data analysis, prepared figures, and wrote the manuscript. T.S. designed the research, supervised the study, reviewed data and results, wrote the manuscript, and secured funding.

