## Supplementary Information for "Multiscale chromatin modeling of chromosome X structural changes upon inactivation highlights the differential regulatory mechanism of *Xist*"

**Supplementary Note 1.**

#### Additional details on the nucleosome-resolution model

- **Energy terms:**

  The total energy associated with the nucleosome-resolution model (Eq. 1 in the main text) incorporates six types of interactions that together capture the mechanical, electrostatic, and epigenetically modulated behavior of chromatin fibers. These terms include stretching, bending, twisting, electrostatics, excluded volume, and methylation-mediated contacts. Below, we describe the functional form and physical interpretation of each contribution.

  – Stretching: The stretching energy between bonded bead pairs is modeled using a harmonic potential:

  $$E_{S_i} = \frac{h_i}{2}(l_i - l_0^i)^2.$$

  Here, $h_i$ is the stretching stiffness associated with the $i^{\text{th}}$ bond, $l_i$ is the instantaneous bond length, and $l_0^i$ is the corresponding equilibrium bond length. Both $h_i$ and $l_0^i$ depend on the specific type of bonded pair (e.g., linker DNA, nucleosome core, linker histone, histone tails).

  – Bending: Local bending rigidity is incorporated through a harmonic potential applied to consecutive triplets of beads:

  $$E_{B_i} = \frac{g_i}{2}(\beta_i - \beta_0^i)^2.$$

  Here, $g_i$ is the bending stiffness, $\beta_i$ is the instantaneous bond angle, and $\beta_0^i$ is its equilibrium value. As with the stretching term, the constants depend on the identity of the beads forming the angle and encode the characteristic flexibility of DNA and nucleosome linkers.

*Electronic address: `Correspondingauthor:`

– Twisting: Torsional rigidity is included for linker DNA segments and nucleosome cores:

$$E_{T_i} = \frac{s}{2l_0}(\phi_i - \phi_0^i)^2.$$

Here, $s$ is the torsional stiffness, $l_0$ is the equilibrium bond length, $\phi_i$ is the torsional angle for the $i^{\text{th}}$ triplet, and $\phi_0^i$ is its equilibrium value. This term ensures proper representation of DNA helicity and rotational constraints around nucleosomes.

– Electrostatics: Long-range electrostatic interactions between charged beads are computed using the screened Debye-Hückel potential:

$$E_C(q_i, q_j, r_{ij}) = \frac{q_i q_j}{4\pi\epsilon_0\epsilon_r r_{ij}} e^{-\kappa r_{ij}}.$$

Here, $q_i$ and $q_j$ are the bead charges, $r_{ij}$ is their separation, $\epsilon_0$ is the vacuum permittivity, $\epsilon_r$ is the relative dielectric constant, and $\kappa$ is the inverse Debye length. Electrostatic interactions are calculated for all charged bead pairs, not restricted by connectivity constraints.

– Excluded volume: Steric repulsion between beads is represented using a truncated Lennard-Jones potential:

$$E_{\text{LJ}}(r_{ij}, \epsilon_{ij}, \sigma_{ij}) = 4\epsilon_{ij}\left[\left(\frac{\sigma_{ij}}{r_{ij}}\right)^{12} - \left(\frac{\sigma_{ij}}{r_{ij}}\right)^{6}\right] \quad \text{for } r_{ij} < r_c.$$

Here, $\epsilon_{ij}$ is the interaction depth, $\sigma_{ij}$ is the effective bead diameter, and $r_{ij}$ is the inter-bead distance. This potential prevents steric overlap while allowing appropriate short-range bead repulsion.

– Methylation contacts: To incorporate the experimentally observed tendency of methylated nucleosomes to form long-range interactions, we include additional harmonic contacts between methylated sites:

$$E_{\text{methyl}} = \frac{h_m}{2}(r_{ij} - r_0)^2.$$

Here, $h_m$ is the spring constant associated with methylation-mediated attraction, $r_0$ is the equilibrium distance, and $r_{ij}$ is the separation between the interacting nucleosomes. These interaction pairs are predicted by a machine-learning model trained on ChIP-seq and Hi-C datasets in methylated and unmethylated conditions (see below), enabling data-driven incorporation of methylation-dependent chromatin folding.

- **Prediction of methylation-associated contacts:**

To identify long-range contacts associated with H3K27me3 within the 90kb X-inactivation center (XIC) region, we used the machine-learning (ML) framework developed by Li et al. [1], which predicts methylation-dependent Hi-C contacts directly from one-dimensional ChIP-seq profiles. Below, we briefly summarize the original approach and describe how the pretrained model was applied in our study.

Summary of the original algorithm:

– Data preparation: Two types of experimental datasets were used as input. (i) ChIP-seq tracks for H3K9me3 or H3K27me3, and (ii) Hi-C datasets generated with and without histone methylation. First, these datasets were standardized to a common resolution. The resulting vector was normalized to zero mean and unit variance, ensuring that methylation intensities lie on a comparable scale across datasets. To construct the methylation-dependent target matrix, the difference between Hi-C maps with and without methylation was computed.

– Feature generation: For each genomic pair $(i, j)$, three features were constructed: the genomic distance $|i - j|$, the methylation intensity at locus $i$, and the methylation intensity at locus $j$. Feature scaling was performed to ensure zero mean and unit variance.

– Model training and evaluation: These features serve as input to a `RandomForestClassifier` (scikit-learn), trained to predict whether each pair corresponds to a methylation-associated long-range contact. Performance was evaluated by comparing predicted methylation-associated contacts against the experimental on/off labels in both the training and testing sets.

Application to the *Xic* simulations

In this work, we did not retrain the machine-learning model; instead, we applied the pretrained model [1]. H3K27me3 ChIP-seq datasets corresponding to the active (Xa) and inactive (Xi) X chromosomes were extracted for the 90kb genomic region in *Xic*. To ensure consistent scaling between the two alleles, we performed joint normalization: the mean and standard deviation were computed from the combined Xa+Xi ChIP-seq dataset, and these parameters were then used to normalize the data for each allele. For each allele, the normalized ChIP-seq profiles were used as input to the pretrained model, which produced predictions of methylation-associated long-range contacts for both Xi and Xa within the simulated *Xic* region.

##### Additional details on the polymer model

- **Cohesin binding and dissociation:** We use kinetic rate events to model the binding/dissociation of cohesin units [2]. When a binding event occurs, cohesin attaches at a randomly selected site where two adjacent beads are unoccupied by CTCF or other cohesins. To maintain realistic steady-state cohesin occupancy, binding and dissociation rates depend on the number of bound cohesins ($N_c$) and available free sites ($N_f$):

$$\begin{aligned} r_+ &= r^0_\pm \quad \frac{1}{\eta + \exp\big(\mu[2N_c - N_f - A_0]\big)} \\ r_- &= r^0_\pm \quad \frac{1}{1 + \exp\big(-\mu[2N_c - N_f - A_0]\big)} \end{aligned} \tag{1}$$

  Here,$A_0 = 2N_c^0 - N_f^0$ is an asymmetry parameter defined by steady-state values $N_c^0$ and $N_f^0$. The parameter $\eta = \frac{N_f^0}{N_c^0} - 1$ ensures detailed balance (steady-state condition $N_f^0\ r_+ = 2N_c^0\ r_-$, while $\mu$ controls fluctuations around equilibrium and $r^0_\pm$ sets the attempt rate.

- **Loop extrusion dynamics:** Once bound, each cohesin's two anchors slide independently in opposite directions at rate $r_s = 2N_c r_s^0$, where $r_s^0$ is the intrinsic sliding rate per anchor. Sliding continues until an anchor encounters oppositely oriented bound CTCF barrier or another cohesin, at which point that anchor pauses while the opposite anchor may continue extending the loop. This asymmetric stalling reproduces the unidirectional blocking effect of convergent CTCF sites observed experimentally.

#### Supplementary Note 2. Additional details on quantities measured

- **Contact probability:**

  The contact probability quantifies the likelihood of physical interaction between any two nucleosome cores along the chromatin fiber. For the nucleosome-resolution model, contacts between nucleosomes are mapped to their corresponding genomic bins (of size 200 bp) based on their positions along the sequence. The contact probability between two nucleosomes belonging to genomic bins $\alpha$ and $\beta$ is then obtained by averaging over all configurations and simulation trajectories as follows:

$$P_{\alpha\beta} = \left\langle \frac{1}{n_\alpha n_\beta} \sum_{i\in\alpha} \sum_{j\in\beta} H(r_{\text{cut}} - r_{ij}) \right\rangle, \tag{2}$$

where: $r_{ij}$ is the distance between nucleosomes $i$ and $j$. $H(r)$ is the Heaviside step function, and $r_{\text{cut}} = 25$nm defines the contact threshold. $n_\alpha$ and $n_\beta$ are the numbers of nucleosomes within bins $\alpha$ and $\beta$, respectively. Angular brackets denote the average over all configurations and trajectories. This normalization ensures that $P_{\alpha\beta}$ reflects the average probability that a nucleosome from bin $\alpha$ is in contact with any other nucleosome from bin $\beta$.

For coarse-grained polymer simulation, we use a similar method to compute the contact probability between monomers of size 2kb. The contact probability between beads $i$ and $j$ is given by $P_{ij} = \langle H(r_{\text{cut}} - r_{ij}) \rangle$, where $r_{ij}$ is the 3D distance between the CG beads, $r_{\text{cut}} = 1.5\sigma$ is the cut-off distance, and the angular brackets denote the average over all configurations.

- **Radius of gyration:**

We compute the radius of gyration $R_g$ to quantify the compaction of chromatin polymer. For the nucleosome-resolution model, we use nucleosome cores alone to compute $R_g$ as the mass of linker beads is much less than that of the nucleosome core. The average radius of gyration of the polymer is given by

$$R_g = \sqrt{\left\langle \frac{1}{N_c} \sum_{i=1}^{N_c} |\mathbf{r}_i - \mathbf{r}_{\text{com}}|^2 \right\rangle}. \tag{3}$$

Here, $N_c$ is the number of nucleosome cores in the corresponding trajectory, $\mathbf{r}_i$ is the position vector of the $i^{th}$ nucleosome core, and $\mathbf{r}_{\text{com}}$ is the center of mass position vector ($\mathbf{r}_{\text{com}} = 1/N_c \sum_i \mathbf{r}_i$.).

For the polymer model, we compute the $R_g$ using a similar method as above for the corresponding $N$ polymer beads.

- **Nucleosome clutch analysis:**

A nucleosome clutch is defined as a spatial cluster of nucleosomes that are mutually in contact. Specifically, two nucleosomes are considered part of the same clutch if their centers lie within a cutoff distance $r_{\text{cut}} = 20$nm. Clutches are identified by assigning a unique cluster index to each connected group of nucleosomes that satisfy this distance criterion. The clutch size corresponds to the number of nucleosomes in a given cluster.

For locus-specific clutch distribution, we consider a clutch part of the locus if more than 50% of nucleosomes in that clutch are with the corresponding gene locus.

- **Stratum-adjusted correlation coefficient (SCC):**

We use the stratum-adjusted correlation coefficient to quantify the reproducibility of the contact maps, thereby removing biases that arise from the dependence of contact probability on genomic separation [3]. The SCC algorithm stratifies the data by genomic distance, thereby removing the distance dependence when comparing experimental and simulated contact maps.

Before computing the SCC, we smooth each contact map using a two-dimensional mean filter to eliminate experimental noise. Specifically, each contact probability value $P_{ij}$ is replaced by the mean over a square neighborhood of side length $2h + 1$, centered at $P_{ij}$. We use $h = 10$ for the smoothing.

After smoothing, the data are divided into $K$ strata based on the contour distance $|i - j|$, where $K = 100$. For each stratum, we compute the Pearson correlation coefficient between the experimental and simulated contact maps using the log-transformed values $\log(P_{ij})$. The SCC is obtained as the weighted sum of these stratum-specific correlation coefficients [3].

- **3D distance map:**

  We generate 3D distance map by computing the average 3D distance between nucleosome cores belonging to two genomic bins along the chromatin fiber. For the nucleosome-resolution model, distances between individual nucleosomes are mapped to their corresponding genomic bins (of size 200 bp) based on their genomic positions. The average 3D distance between nucleosomes belonging to genomic bins $\alpha$ and $\beta$ is computed by averaging over all nucleosome pairs, configurations, and simulation trajectories as follows:

$$D_{\alpha\beta} = \left\langle \frac{1}{n_\alpha n_\beta} \sum_{i\in\alpha} \sum_{j\in\beta} r_{ij} \right\rangle, \tag{4}$$

  where $r_{ij}$ is the 3D distance between nucleosomes $i$ and $j$, and $n_\alpha$ and $n_\beta$ denote the numbers of nucleosomes within genomic bins $\alpha$ and $\beta$, respectively. Angular brackets indicate averaging over all configurations and trajectories. This normalization ensures that $D_{\alpha\beta}$ represents the mean 3D distance between a nucleosome from bin $\alpha$ and a nucleosome from bin $\beta$.

#### Supplementary Tables

| Gene/Sequence | Type | Coordinates (mm10) | |
|---|---|---|---|
| | | start | end |
| *Xist* | Non-coding gene | 103460366 | 103483254 |
| *Xite* | Regulatory element | 103426201 | 103438709 |
| *Jpx* | Non-coding gene | 103493558 | 103506425 |
| *Tsix* | Non-coding gene | 103431517 | 103484957 |
| *Rnf12* | Coding gene | 103962494 | 103966656 |
| *Linx* | Non-coding gene | 103210000 | 103313000 |
| *Ftx* | Non-coding gene | 103569505 | 103623754 |
| *Chic1* | Non-coding gene | 103356476 | 103396118 |

Supplementary Table 1: Genomic locations of relevant genes and regulatory sequences in the *Xic* region

**Supplementary Figures**

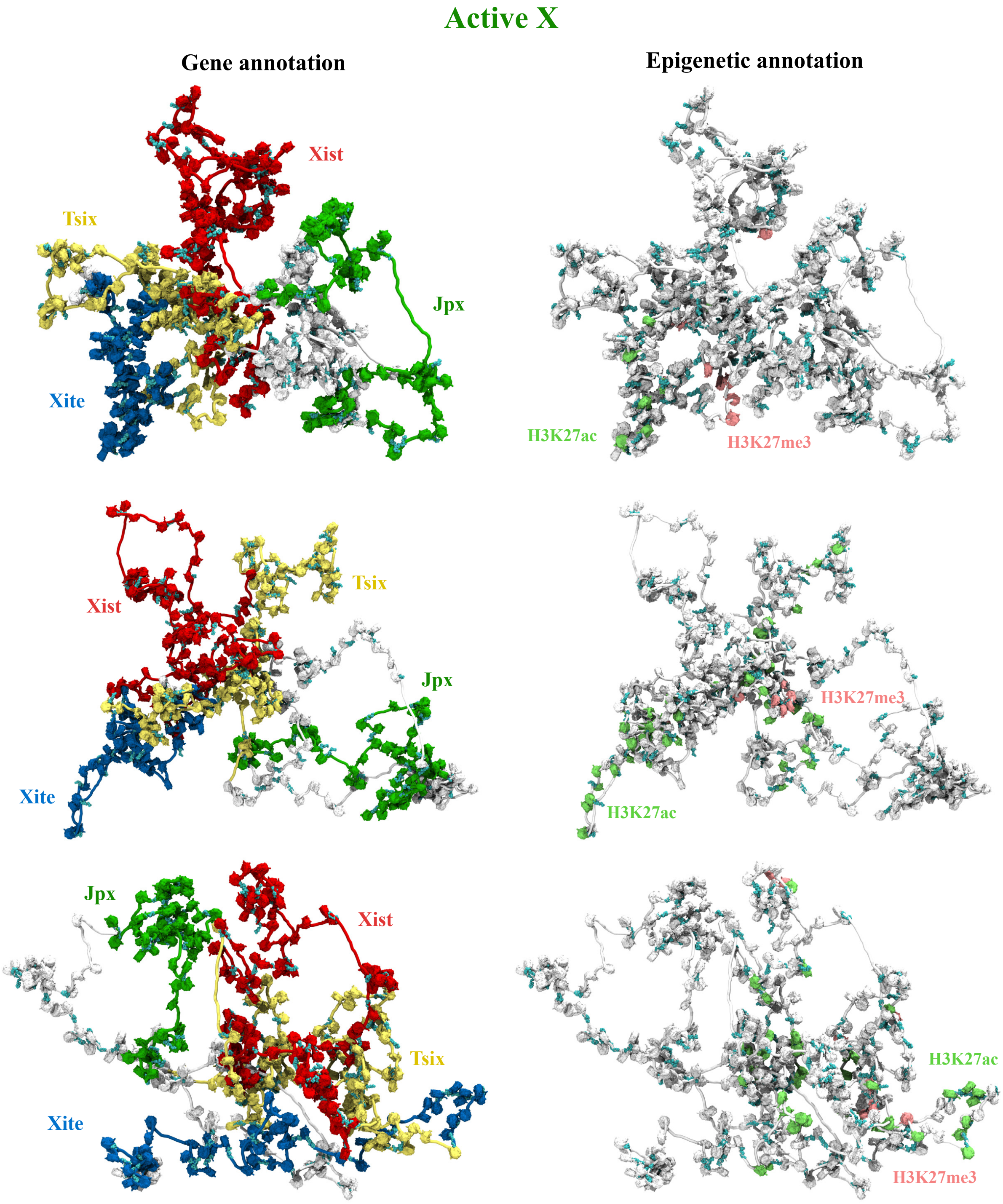


Supplementary Fig. 1: Representative 3D conformations of the 90 kb *Xic* region from the nucleosome-resolution model are shown for the Xa. The left panels are annotated by gene locations, whereas the right panels show the same conformations annotated by epigenetic marks.

### Inactive X

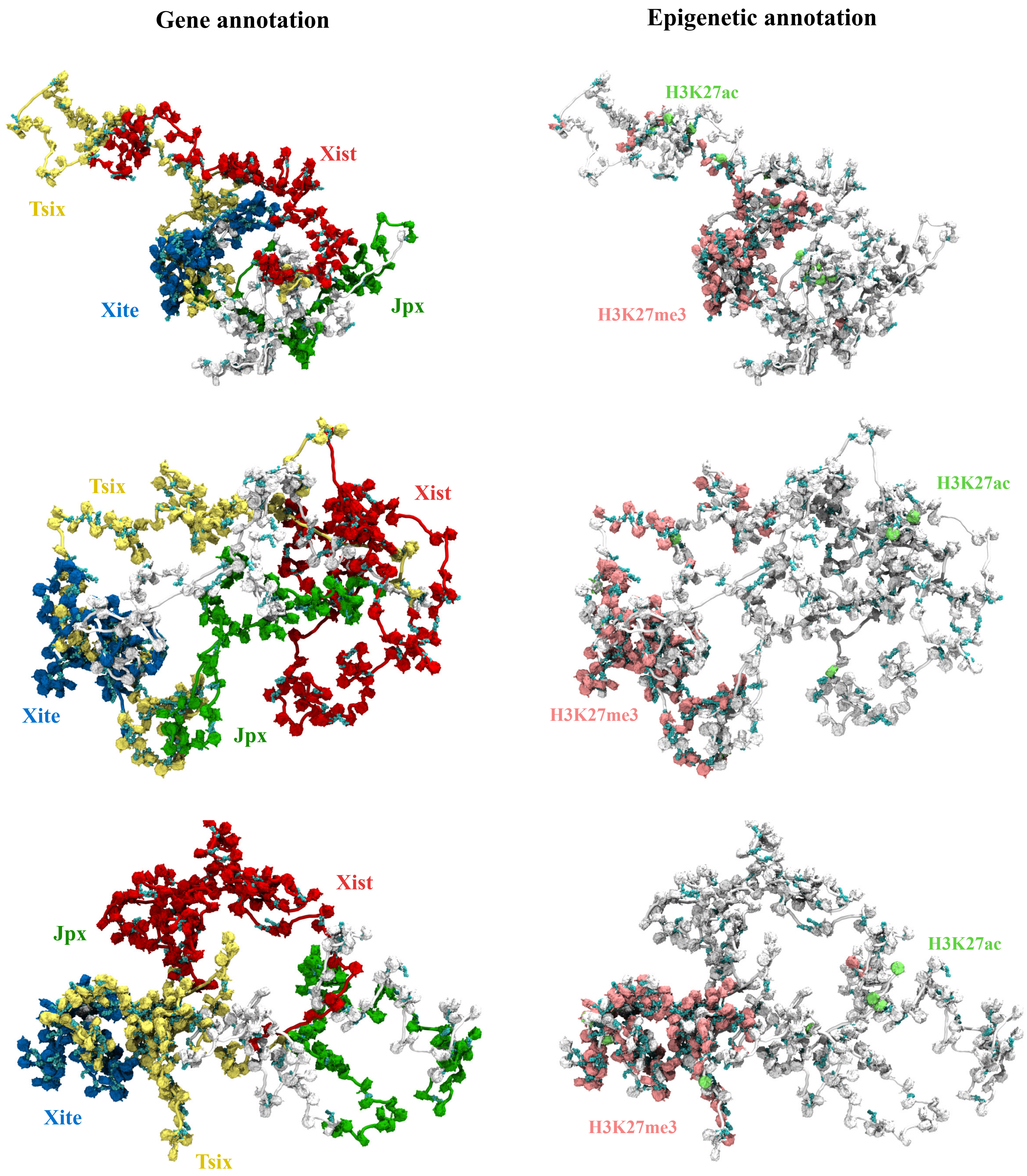


Supplementary Fig. 2: Representative 3D conformations of the 90 kb *Xic* region from the nucleosome-resolution model are shown for the Xi. The left panels are annotated by gene locations, whereas the right panels show the same conformations annotated by epigenetic marks.

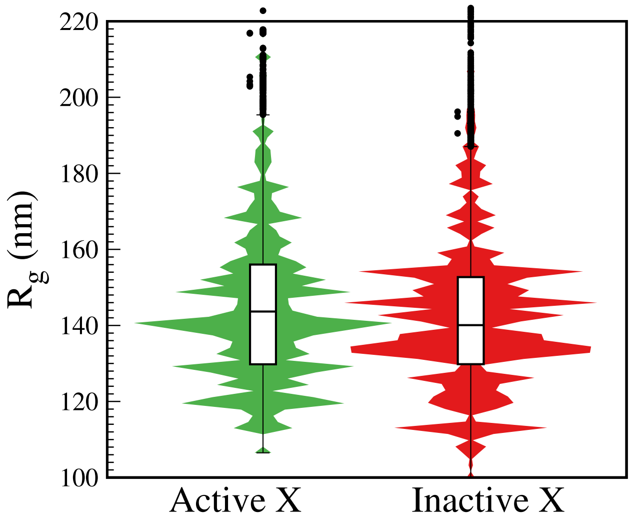


Supplementary Fig. 3: Radius of gyration of the 90kb *Xic* region from nucleosome-resolution model is compared for Xa and Xi using a violin plot.

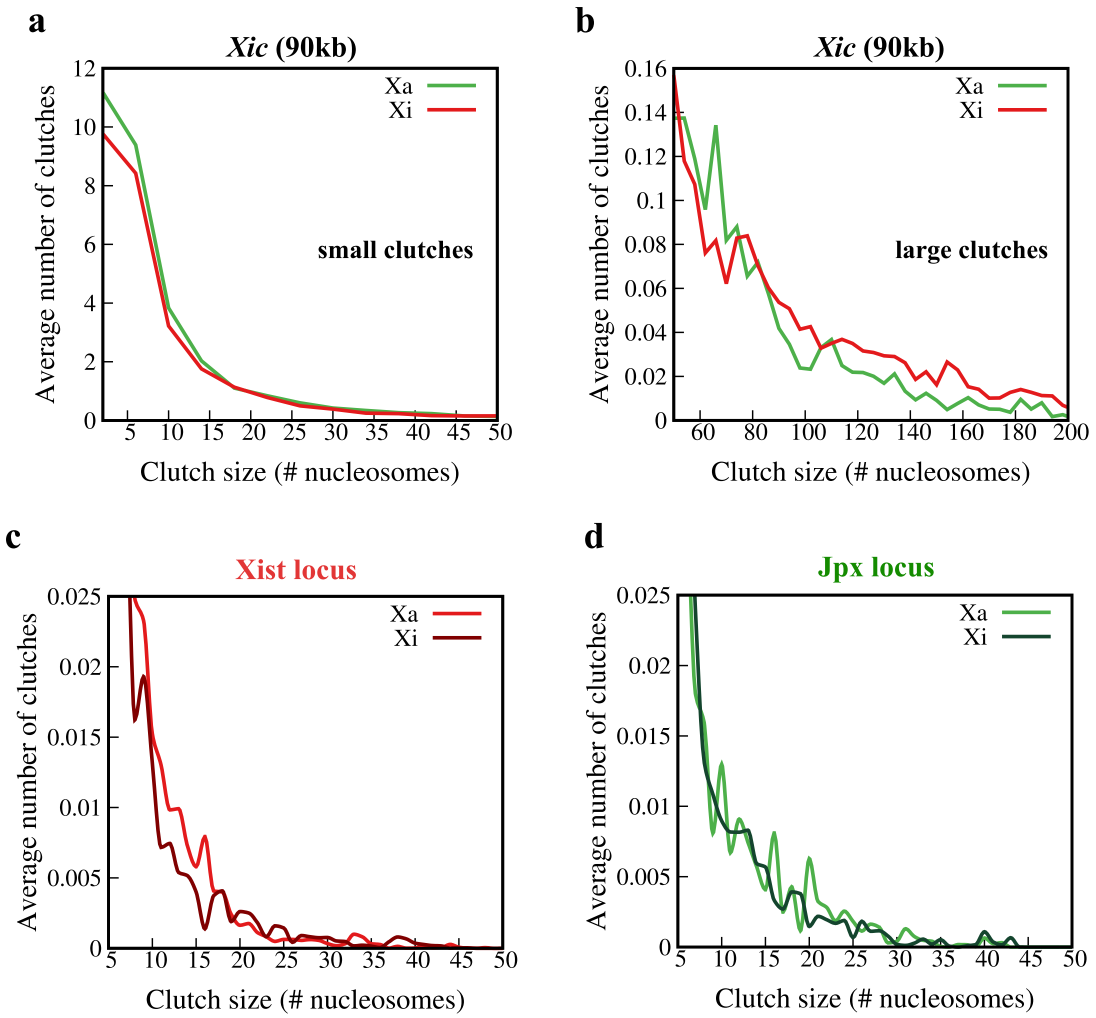


Supplementary Fig. 4: **a-b** Distribution of nucleosome clutch sizes in the 90 kb region for Xa and Xi, shown separately for (**a**) small clutches (fewer than 40 nucleosomes) and (**b**) large clutches (more than 40 nucleosomes). **c-d** Distribution of nucleosome clutch sizes for Xa and Xi at specific loci, shown separately for (**c**) the *Xist* locus and (**d**) the *Jpx* locus.

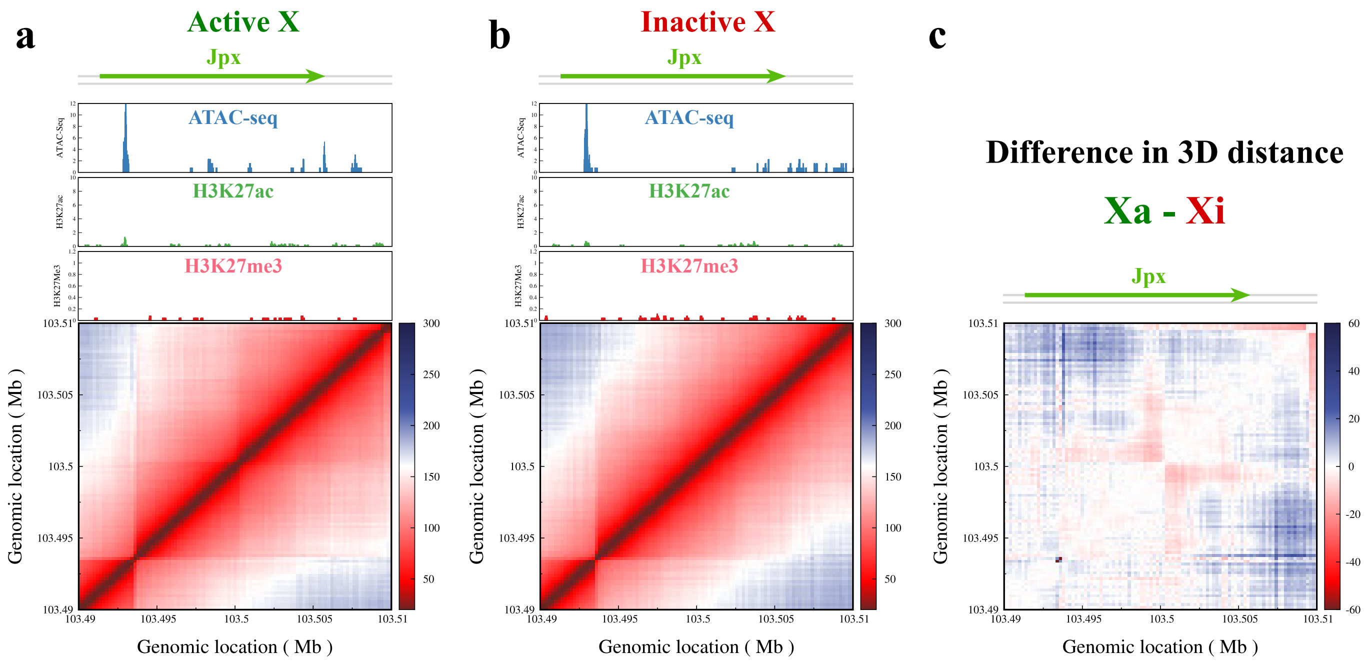


Supplementary Fig. 5: **a-b** 3D distance maps of the *Jpx* locus are plotted for active and inactive X. The genes and regulatory sequences are annotated on top. The corresponding ATAC-seq peaks, denoting DNA accessibility, the active histone mark H3K27ac, and the inactive histone mark H3K27me3, are plotted on top of the distance maps. **c** The difference in average 3D distance (Xi minus Xa) between nucleosomes is plotted as a heatmap with negative values shown in red and positive values shown in blue.

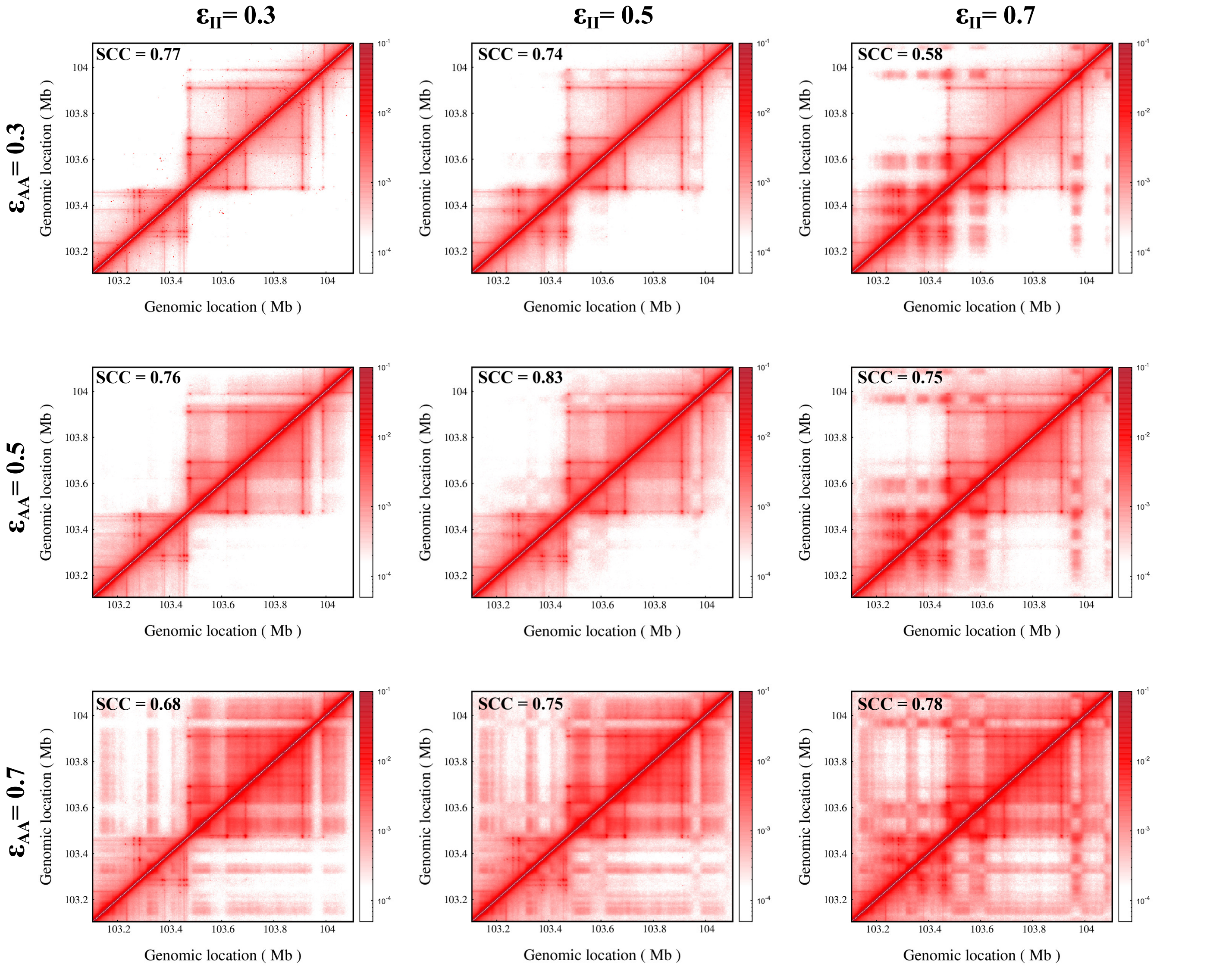


Supplementary Fig. 6: Contact maps of the 1Mb region from polymer simulations are plotted for different values of self-interactions between active type A ($\epsilon_{AA}$) and inactive type I beads ($\epsilon_{II}$). The interaction between type A and type I beads is fixed $\epsilon_{AI} = 0.2$. The stratum-adjusted correlation coefficient between the simulation and experimental contact maps is shown on the top left of the corresponding simulation maps.
